# Multi-Pathway DNA Double-Strand Break Repair Reporters Reveal Extensive Cross-Talk Between End-Joining, Single Strand Annealing, and Homologous Recombination

**DOI:** 10.1101/2021.12.22.473809

**Authors:** Bert van de Kooij, Alex Kruswick, Haico van Attikum, Michael B. Yaffe

## Abstract

DNA double-strand breaks (DSB) are repaired by multiple distinct pathways, with outcomes ranging from error-free repair to extensive mutagenesis and genomic loss. Repair pathway cross-talk and compensation within the DSB-repair network is incompletely understood, despite its importance for genomic stability, oncogenesis, and the outcome of genome editing by CRISPR/Cas9. To address this, we constructed and validated three fluorescent Cas9-based reporters, named DSB-Spectrum, that simultaneously quantify the contribution of multiple distinct pathways to repair of a DSB. These reporters distinguish between DSB-repair by error-free canonical non-homologous end-joining (c-NHEJ) versus homologous recombination (HR; reporter 1), mutagenic repair versus HR (reporter 2), and mutagenic end-joining versus single strand annealing (SSA) versus HR (reporter 3). Using these reporters, we show that inhibition of the essential c-NHEJ factor DNA-PKcs not only increases repair by HR, but also results in a substantial increase in mutagenic repair by SSA. We show that SSA-mediated repair of Cas9-generated DSBs can occur between Alu elements at endogenous genomic loci, and is enhanced by inhibition of DNA-PKcs. Finally, we demonstrate that the short-range end-resection factors CtIP and Mre11 promote both SSA and HR, whereas the long-range end-resection factors DNA2 and Exo1 promote SSA, but reduce HR, when both pathways compete for the same substrate. These new Cas9-based DSB-Spectrum reporters facilitate the rapid and comprehensive analysis of repair pathway crosstalk and DSB-repair outcome.

## Introduction

Double-strand DNA breaks (DSBs) are severe genotoxic lesions that need to be correctly repaired to prevent mutations, loss of genomic information or devastating chromosomal rearrangements. DSBs result from exogenous sources like environmental radiation or anti-cancer chemotherapeutics, or from endogenous sources like replication stress or endogenous nucleases that function during meiosis and immune cell maturation^1^. Furthermore, site-specific DSB-formation by Cas9 or related endonucleases is central to CRISPR-based genome editing, which has become an essential technology in biomedical research, and holds great promise to cure disease in gene therapy applications^2–4^.

Mammalian cells are equipped with multiple pathways to repair DSBs^5^. In most cells, the dominant DSB-repair pathway is canonical Non-Homologous End-Joining (c-NHEJ)^5^. DSB-repair by c-NHEJ is mostly accurate, but can also introduce small insertions and deletions (InDels) at the DSB junction caused by minor editing of the DSB-ends prior to ligation^6^. As an alternative to c-NHEJ, DSBs can be repaired by Homologous Recombination (HR). This starts with end-resection of the DSB-ends to generate single-strand 3’ overhangs that subsequently invade homologous DNA, usually the sister chromatid, which functions as a template that is copied in the downstream repair steps^7^. HR is therefore considered an error-free pathway. However, DNA fragments that share homology with the DSB-site but carry mutations or even large insertions can also be copied during repair by HR. This is utilized for genome editing purposes by co-delivery of a repair template with the desired mutant sequence. Compared to genome editing by c-NHEJ, this expands the editing possibilities and allows additional control over the editing outcome, but this approach is hampered by the low frequency of HR compared to c-NHEJ^5^. Many strategies to increase the ratio of HR to c-NHEJ have been developed, but c-NHEJ is surprisingly robust and remains the dominant repair pathway even when applying these HR-promoting methods^8^.

In addition to c-NHEJ and HR, DSBs can be repaired by alternative End-Joining (a-EJ), or Single-Strand Annealing (SSA)^9,10^. Both pathways require end-resection to expose regions of homology on the same broken DNA molecule and adjacent to the DSB. Annealing of these homologous regions is followed by removal of non-homologous resected DNA, polymerase-mediated fill-in and ligation. Compared to a-EJ, SSA requires larger regions of homology, and is more tolerant to large stretches of DNA separating the homologous regions^11^. Both a-EJ and SSA are inherently mutagenic, and SSA, in particular, can result in loss of multiple kilobases of genetic information^12^. How frequently these pathways act on genomic DSBs is not known, but recent studies suggested that a substantial fraction of Cas9-generated DSBs may be repaired by a-EJ^13,14^. The contribution of SSA to repair of Cas9 DSBs, however, has not been thoroughly addressed.

These mechanistically distinct DSB-repair pathways do not function independently, but act in a network that is tightly regulated to optimally preserve the integrity of the genome. Deregulation of the DSB-repair network reduces genomic stability and can have severe pathogenic consequences, most notably cancer development^15^. Intense research efforts are slowly revealing the organization of this network, and have shown that end-resection dependent repair by HR is promoted by BRCA1, whereas the 53BP1-Shieldin complex inhibits end-resection and therefore directs DSB-repair towards c-NHEJ^16–21^. Nevertheless, many nodes and edges of the DSB-repair network remain to be revealed. For example, whereas it has generally been well-described how inhibiting c-NHEJ, or promoting resection of DSB ends, affects repair by HR, most studies have not reported how this affects repair by other end-resection dependent pathways like a-EJ, or SSA (reviewed in Yeh *et al*.)^8^. Moreover, how DSB-repair pathway choice between a-EJ, SSA and HR is regulated after the initiation of end-resection is not well described. These are import questions to answer since there is a strong interest in controlled manipulation of DSB-repair pathway choice, to improve anti-cancer therapy, and more recently to direct genome editing outcome^8,22^.

To facilitate network-level analysis of DSB-repair activity, we developed three variants of a novel genomic Cas9-based DSB-repair reporter construct that we named DSB-Spectrum. Reporter constructs that quantify activity of DSB-repair pathways, such as DR-GFP, which quantifies the frequency of HR, have greatly contributed to DSB-repair research^23^. Multiple elegant variations of DR-GFP-type reporters have been generated that quantify repair by c-NHEJ, a-EJ, or SSA^24–26^, and in some cases a combination of these pathways^27,28^. Using these published reporter systems as a basis, we generated the DSB-Spectrum variants, which are multi-pathway reporters that allow for simultaneous quantification of the frequency of DSB-repair by end-joining, SSA and HR.

These DSB-Spectrum reporters display high frequencies of DSB-repair through all pathways, function as single constructs without the requirement for ectopic HR-donors, and are activated by Cas9-induced DSB generation, allowing for direct translation of the results to CRISPR-strategies. We show that DSB-Spectrum efficiently and correctly reveals DSB-repair pathway crosstalk between NHEJ, SSA and HR. We demonstrate that SSA can strongly contribute to DSB-repair and is significantly promoted by inhibition of DNA-PKcs. We show that repetitive elements in the human genome can drive SSA of Cas9-induced DSBs in endogenous genomic loci. Finally, we demonstrate that SSA, but not necessarily HR, is promoted by the long-rang end-resection factors Exo1 and DNA2, thus providing a potential mechanism to direct DSB-repair away from SSA towards HR. Our studies show that DSB-Spectrum captures the complexity of the DSB-repair network and detects DSB-repair phenotypes that can easily be missed by commonly used sequencing approaches that analyze the DSB-repair junction, or by studies that focus on individual repair pathways.

## Results

### DSB-Spectrum_V1 is a multi-pathway DSB-repair reporter that simultaneously quantifies c-NHEJ and HR

To study the interaction between the various DSB-repair pathways, we sought to design a multi-pathway reporter in which repair of a single site-specific DSB by each individual pathway would result in a unique, pathway-specific expression pattern of fluorescent proteins (Fig. 1A). Such a reporter would reveal, on an individual cell basis, which pathway was used to repair the DSB. Within a multicellular population, such a reporter system could be used to simultaneously quantify the frequency of repair by multiple DSB-repair pathways using flow cytometry as a readout.

**Figure 1.**
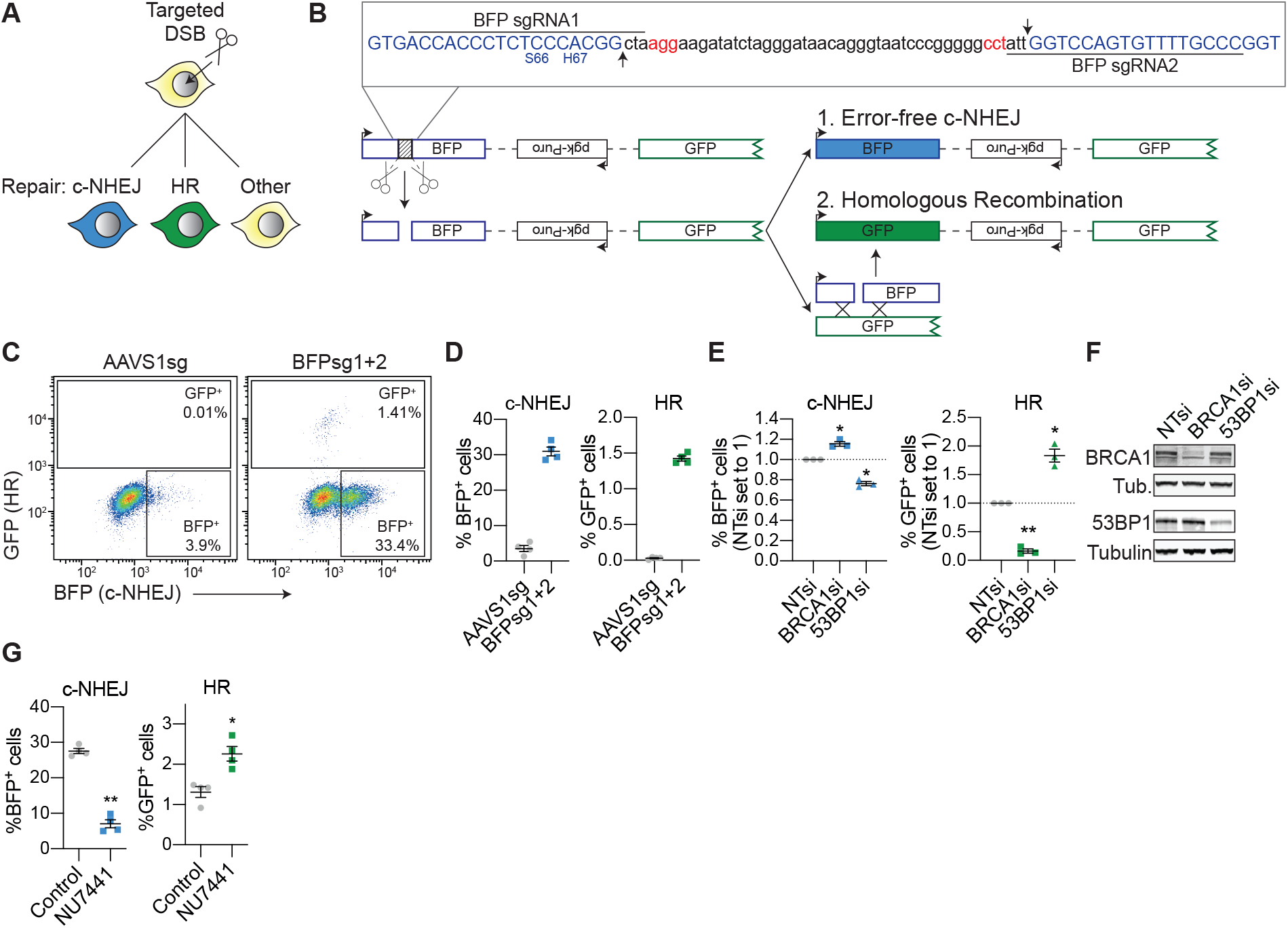
DSB-Spectrum_V1 is a reporter for both c-NHEJ and HR. **(A)** Cartoon depicting potential outcomes of a multi-pathway reporter cell-line designed to quantify DSB-repair by error-free c-NHEJ and HR. **(B)** Diagramatic representation of the genomic DSB-repair reporter construct DSB-Spectrum_V1. Expanded region shows the DNA sequence targeted by Cas9. The sequence of the BFP cDNA is displayed in blue, the PAM sequences of the sgRNA target sites are displayed in red. Arrows in the inset and scissors in the cartoon indicate the Cas9 cut sites. Ligation of the distal DSB-ends by error-free c-NHEJ will remove the spacer sequence and restore the BFP gene. **(C and D)** DSB-Spectrum_V1 was integrated into the genome of HEK 293T cells by lentiviral infection, followed by expansion of a single cell clone. The resulting DSB-Spectrum_V1 cell-line was transfected with Cas9 and an sgRNA targeting a control locus (AAVS1) or BFP, and analyzed by flow cytometry at 72h after transfection. Panel C shows representative flow plots, and panel D shows the quantification of multiple experiments (n=4; mean ± SEM). **(E)** DSB-Spectrum_V1 cells were transfected with indicated siRNAs (NTsi=Non-Targeting control), followed by transfection with Cas9 and an sgRNA targeting a control locus or BFP. At 72h after Cas9 transfection cells were analyzed by flow cytometry. Percentages of each fluorescent population in the BFPsg-transfected cells were corrected for the background percentages seen in the AAVS1sg-transfected cells. Depicted is the ratio of background-corrected percentages for each fluorescent population to that of the NTsi control (n=3; mean ± SEM; **p≤0.01; *p≤0.05). **(F)** Western blot of lysates from cells analyzed in panel E. Tubulin is used as loading control. **(G)** As in panel D, but including treatment with NU7441 (2 µM), and background-corrected as described in panel E (n=4; mean±SEM; **p≤ 0.01; *p≤0.05).

To accomplish this, we first modified and combined two existing reporter systems to create a multi-pathway reporter construct designed to detect distinct repair products created by either c-NHEJ or HR (Fig. 1A).^23,25^ Figure 1B shows this reporter, which we named DSB-Spectrum_V1, and which consists of a CMV promoter followed by a modified gene encoding Blue Fluorescent Protein (BFP), a 2.6 kb intervening region, and a truncated gene encoding part of Green Fluorescent Protein (GFP) and lacking a promoter. Hence, no GFP is expressed from the reporter. No functional BFP is produced either, because the BFP gene contains a 46 bp spacer insert separating the gene at the triplet encoding for Gly-68, immediately adjacent to the critical amino acid, His-67, responsible for blue fluorescence. This spacer sequence can be excised by targeting Cas9 to the edges using sequence-specific guide RNAs, leaving behind a blunt ended DSB. Ligation of the DSB through error-free c-NHEJ would restore the intact BFP sequence, and result in BFP expression. Alternatively, the DSB can be repaired by Homologous Recombination (HR) using the truncated GFP gene as a repair template, since it shares high sequence homology with the BFP gene (Fig. S1A, B). Repair by HR will thus result in gene conversion and expression of GFP (Fig. 1B). Error-free c-NHEJ and HR can therefore be clearly distinguished from mutagenic repair pathways, because these latter types of repair would not result in BFP or GFP expression (Fig. 1A).

A single copy of DSB-Spectrum_V1 was integrated into the genome of HEK 293T cells by lentiviral delivery, and a single cell clone expanded to obtain a homogeneous DSB-Spectrum_V1 cell-line. For this DSB-Spectrum_V1 clone, and for other DSB-Spectrum clones used in this manuscript, Splinkerette PCR was used to map the genomic integration site of the reporter^29^. For all clones, only one integration site was retrieved, consistent with single integration. Furthermore, for all clones, the sites of integration of the reporter construct were either within large introns or intergenic regions of genes that are not associated with canonical modes of DNA repair (Fig. S1C).

Next, these cells were transfected with Cas9 cDNA and either an sgRNA targeting a control genomic locus (AAVS1sg), or two sgRNAs targeting the spacer region in the reporter (BFPsg1+2), followed by flow cytometric analysis 72 hours post transfection. As shown in figures 1C and D, distinct BFP^+^ and GFP^+^ populations could be detected specifically upon targeting Cas9 to the reporter. Approximately 30% of cells were BFP^+^, while ∼1.5% of cells were GFP^+^, consistent with a minority of DSBs being repaired by HR compared to error-free c-NHEJ (Fig. 1D). To validate that the BFP^+^ and GFP^+^ populations resulted from repair of the Cas9-DSBs by c-NHEJ or HR, respectively, we used RNAi to deplete either BRCA1, an HR-promoting factor, or 53BP1, which is a well-established c-NHEJ-promoting factor. As shown in figures 1E and F, depletion of BRCA1 strongly reduced the frequency of GFP^+^ cells, while it increased the frequency of BFP^+^ cells. The opposite phenotype was observed upon loss of 53BP1 (Fig. 1E, F; see Fig. S6 for uncropped blots). These data are fully consistent with the known functions of 53BP1 and BRCA1 in DSB-repair^5^, and confirm that the BFP^+^ and GFP^+^ populations resulted from end-joining and HR, respectively.

To further validate DSB-Spectrum_V1 as a reporter for c-NHEJ, rather than other types of end-joining, we individually silenced the expression of several core c-NHEJ factors Ku80, DNA-PKcs, XRCC4 or Lig4, or silenced expression of the a-EJ factor Polθ. Depletion of each of the c-NHEJ factors significantly reduced the frequency of BFP^+^ cells, while no such effect was observed after depletion of Polθ, consistent with BFP expression resulting from DSB repair by c-NHEJ rather than by a-EJ (Fig. S1D, E). Next, we assessed the reporter phenotype following treatment with NU7441, a small molecule inhibitor of DNA-PKcs kinase activity^30^. As shown in Figure 1G, NU7441 treatment resulted in a more than three-fold reduction in the frequency of error-free repair by c-NHEJ. It also resulted in a close to two-fold increase in the percentage of GFP^+^ cells, demonstrating that loss of c-NHEJ is compensated for by an increase in HR. Taken together, these results show that DSB-Spectrum_V1 can reveal the interdependence and crosstalk between the c-NHEJ and HR DSB-repair pathways within a single population of cells.

### DSB-Spectrum_V2 quantitatively reports mutagenic end-joining and HR

To generate a complementary reporter system capable of quantifying mutagenic end-joining, rather than error-free c-NHEJ, together with HR, a variant of DSB-Spectrum_V1 was developed (Fig. 2A, B). The resulting DSB-Spectrum_V2 expresses a functional BFP gene under basal conditions, in contrast to the spacer-separated variant encoding non-functional BFP. A single sgRNA can then be used to target Cas9 to the sequence in the BFP gene that differs from GFP, and is essential for its blue fluorescence (Fig. S1A, B). Following generation of a DSB by Cas9, repair by mutagenic end-joining can be monitored by loss of BFP expression while HR repair, as in DSB-Spectrum_V1, can be monitored as gain of GFP expression. Repair of the DSB by error-free c-NHEJ restores BFP expression, but this repair mechanism cannot be quantified with this reporter construct, since a similar phenotype would result from failure of Cas9 cleavage (labelled ‘other’ in Fig 2A).

**Figure 2.**
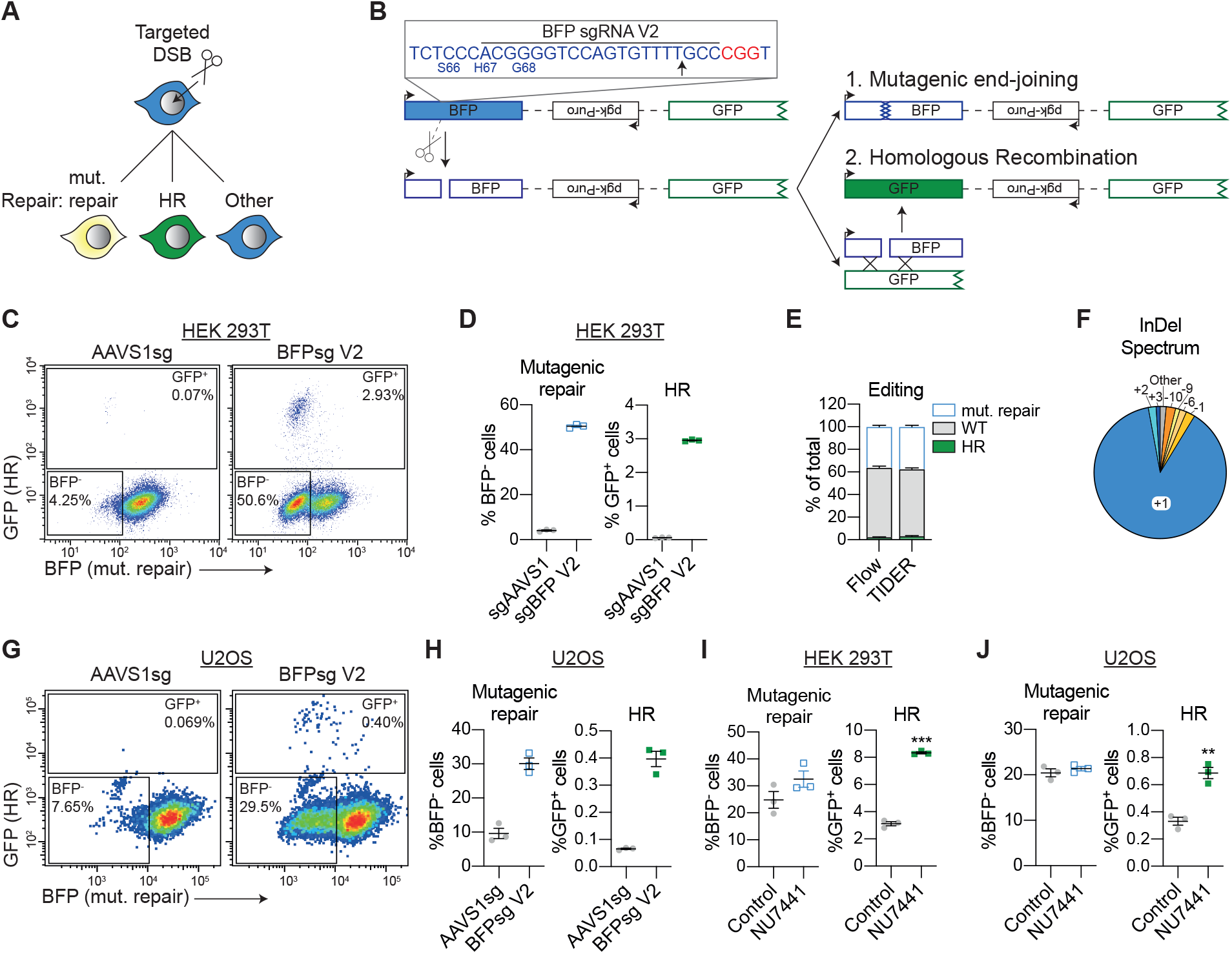
DSB-Spectrum_V2 is a reporter for both mutagenic repair and HR. **(A)** Cartoon depicting potential outcomes of a multi-pathway reporter cell-line designed to quantify DSB-repair by mutagenic end joining and HR. **(B)** Diagramatic representation of the genomic DSB-repair reporter construct DSB-Spectrum_V2. Expanded region shows the DNA sequence targeted by Cas9. The sequence of the BFP cDNA is displayed in blue, the PAM sequence of the sgRNA target site is displayed in red. Arrow in the inset and scissors in the cartoon indicate the Cas9 cut site. **(C and D)** Flow cytometry and quantification of mutagenic repair and HR in DSB-Spectrum_V2 cells (n=3; mean ± SEM), as in figure 1C and D. **(E)** DSB-Spectrum_V2 cells were transfected with Cas9 and an sgRNA targeting either a control locus or BFP. At 48h after Cas9 transfection genome editing was quantified by TIDER analysis of the sequenced target site. At 72h after Cas9 transfection cells were analyzed by flow cytometry (n=3; mean ± SEM). **(F)** The InDel spectrum at the Cas9 target site in DSB-Spectrum_V2, determined by TIDER analysis described in panel E. Pie chart shows the seven most frequently identified InDels. **(G and H)** As in panels C and D, but for U2OS DSB-Spectrum_V2 cells (n=3; mean ± SEM). **(I)** Mutagenic repair and HR was quantified in HEK 293T DSB-Spectrum_V2 cells with or without treatment with NU7441 (2μM; n=3; mean±SEM; ***p≤0.001). **(J)** As in panel I, but for U2OS DSB-Spectrum_V2 cells (n=3; mean±SEM; ***p ≤0.01)

Following integration of a single copy of DSB-Spectrum_V2 into the genome of HEK 293T cells, a clonal cell-line was generated (Fig. S1C). Cells were then transfected with Cas9 together with an sgRNA targeting either a control genomic locus (AAVS1sg), or BFP, and DSB repair outcomes were monitored by flow cytometric analysis. As shown in Figure 2C and D, DSB-Spectrum_V2-expressing cells transfected with the control sgRNA were BFP^+^, consistent with the design of the reporter construct. In contrast, targeting of Cas9 to the reporter resulted in a substantial fraction of the cells losing BFP expression (±50%), while a minor fraction switched from expressing BFP to expressing GFP (almost 3%).

To confirm that loss of BFP expression was the direct result of repair by mutagenic end-joining of the DSB, rather than other epigenetic or genetic mechanisms like promoter silencing or (partial) loss of the DSB-Spectrum reporter, the DNA surrounding the Cas9-cleavage site in DSB-Spectrum_V2 was PCR-amplified, followed by Sanger sequencing of the pool of PCR products. The editing efficiency, as well as the InDel spectrum, was then determined by deconvolving the mixture of sequencing chromatograms by TIDER analysis^31,32^. As shown in figure 2E, the frequencies of wild-type (WT) BFP, mutated BFP, and GFP sequences detected by TIDER analysis reflected the respective frequencies of BFP^+^, BFP^-^ and GFP^+^ cells detected by flow cytometry. Hence, these direct sequencing results verify that loss of BFP was primarily caused by mutagenic end-joining at the DSB site (Fig. 2E). TIDER analysis and sequencing of multiple individual target loci further revealed that the majority of edited repair products contain a +1 thymine insertion at the DSB-junction (Fig. 2F; Fig. S2A), which is typical of editing by c-NHEJ^13,14,33^.

DSB-repair by c-NHEJ can result in gene mutations that fail to disrupt expression of a functional protein product, like silent mutations or in-frame InDels. To validate that none of the BFP^+^ cells contained such phenotypically silent mutagenic end-joining products, we performed a Surveyor nuclease assay^34^. The Cas9 target site was PCR-amplified from the BFP^+^ as well as the BFP^-^ cells, and each PCR product was denatured and re-annealed to itself. If the PCR product was obtained from a mixed pool of wild-type and mutant sequences, the annealing procedure will result in mismatched duplexes that can be digested by the Surveyor nuclease. As expected, Surveyor nuclease digestion products were detected in PCR products from the BFP^-^ population, while no such products were detected in the BFP^+^ population, indicating that these were all WT sequences (Fig. S2C). Thus, the flow cytometry, sequencing and Surveyor nuclease data demonstrate that DSB-Spectrum_V2 is an accurate reporter for both mutagenic end-joining and HR.

To expand the utility of the reporter system we analyzed and validated the performance of DSB-Spectrum_V2 in another widely used cell line, and generated DSB-Spectrum_V2 U2OS cells (Fig. S1C). As observed with the HEK-293T cell-line, both BFP^-^ and GFP^+^ populations were detected in this U2OS clonal cell line specifically after targeting Cas9 to the reporter (Fig. 2G, H). Notably, the gene editing frequencies, in particular the frequency of HR, were lower in U2OS than in HEK 293T cells (Fig. 2D, H), and these editing frequencies were comparable between multiple independent U2OS DSB-Spectrum_V2 clones (Fig. S2D). This suggests that the low editing frequencies are not dependent on the reporter integration site but are a characteristic of U2OS cells. Nonetheless, as all repair populations were present and measurable, these data demonstrate that the DSB-Spectrum_V2 construct consistently functions as a DSB repair reporter in multiple cell types.

The results obtained with DSB-Spectrum_V1 demonstrated that inhibition of DNA-PKcs with the small molecule inhibitor NU7441 reduced c-NHEJ and promoted HR (Fig. 1I). This experiment was repeated with HEK 293T DSB-Spectrum_V2 cells. As expected, a strong increase in HR was observed upon DNA-PKcs inhibition, but surprisingly, this did not result in the expected reduction in BFP loss (Fig. 2I, S2E). Instead, DNA-PKcs inhibition appeared to slightly increase the percentage of BFP^-^ cells, although this failed to reach statistical significance (Fig. 2I). Similar results were obtained in U2OS DSB-Spectrum_V2 cells, where treatment with NU7441 increased HR but failed to reduce mutagenic end-joining (Fig. 2J). These results suggest that, in DSB-Spectrum_V2 cells, the loss of c-NHEJ following NU7441 treatment is compensated for by an alternative type of mutagenic repair that results in BFP loss.

### DNA-PKcs inhibition promotes DSB-repair by SSA

To explore the remaining types of mutagenic repair that might compensate for loss of c-NHEJ in the presence of the DNA-PKcs inhibitor, we investigated whether there might be an increase in repair by a-EJ or SSA. To assess the contribution of a-EJ, an ∼1100 base-pairs (bp) region surrounding the Cas9 cleavage site in DSB-Spectrum_V2 was PCR amplified from both control and NU7441-treated cells, sequenced, and subjected to TIDER analysis. This revealed that NU7441 treatment reduced the frequency of sequences with a 1 bp insertion by ∼15% (Fig. 3A), consistent with inhibition of c-NHEJ by NU7441. We also observed an increased frequency of repair products containing 11 bp or 12 bp deletions (Fig. 3B), which could indicate enhanced repair by a-EJ^33^. However, the total frequency of those a-EJ repair products was consistently less than 2% of the sequenced PCR products, which cannot fully compensate for the reduced frequency of +1 bp repair products resulting from loss of c-NHEJ. Thus, based on this TIDER analysis, it is not evident that a-EJ is responsible for the mutagenic repair causing the loss of BFP expression in the presence of the DNA-PKcs inhibitor.

**Figure 3.**
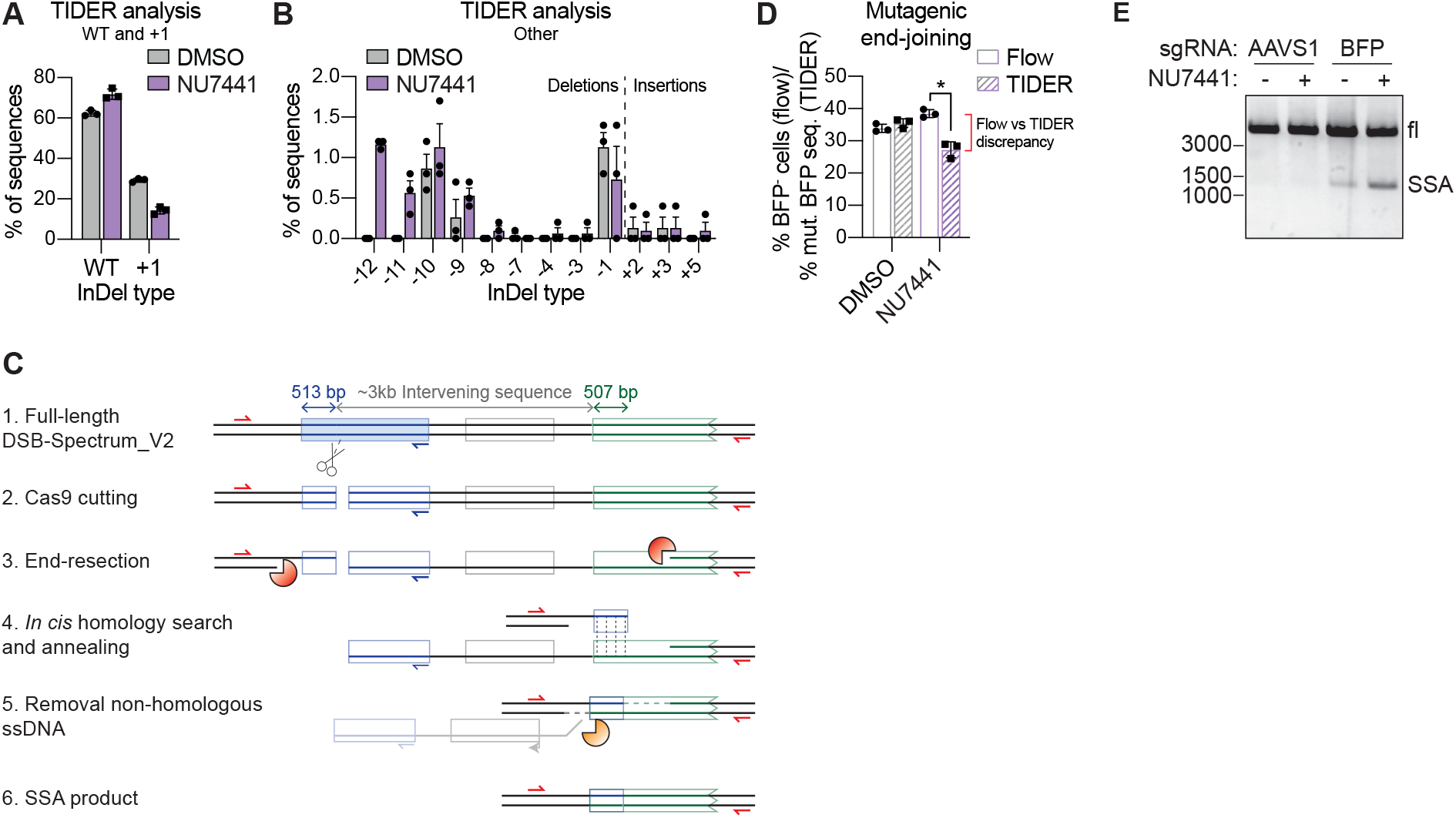
Inhibition of DNA-PKcs promotes DSB-repair by single-strand annealing (A and B) HEK 293T DSB-Spectrum_V2 cells were transfected with Cas9 and an BFP-targeting sgRNA, and subsequently treated with NU7441 (2 µM) or left untreated. At 48h after Cas9 transfection, half of the population was subjected to sequence analysis of the target site using the TIDER algorithm. At 72h after transfection, the remaining half of the population was analyzed by flow cytometry (See panel D). The InDel spectrum is plotted in two graphs to display either the frequency of sequences that are WT or have a +1 insertion (panel A), or the frequency of any of the other detected InDels (panel B; n=3; mean ± SEM; *p≤0.05). **(C)** Schematic representation of repair of the Cas9-induced DSB in DSB-Spectrum_V2 by SSA between the 5’-end of the BFP gene (513bp), located upstream of the DBS, and highly homologous 5’-end of the GFP gene (507bp), located downstream of the DSB. Indicated are the primers (in red) to amplify the SSA repair product, the reverse primer (in blue) to amplify the target site for TIDER analysis, and the nucleases that perform end-resection (orange pacman) or ssDNA flap removal after annealing (yellow pacman). **(D)** The data described in panel A were plotted to show the total frequency of mutant sequences determined by TIDER analysis compared to BFP loss determined by flow cytometry analysis. Note the discrepancy between these analyses following NU7441 treatment (n=3; mean ± SEM; *p≤0.05). **(E)** DSB-Spectrum_V2 cells were transfected with Cas9 and an sgRNA targeting either the AAVS1 locus or BFP, and subsequently treated with NU7441 (2 µM). The genomic DNA was PCR amplified using the primers in red in panel C, and analyzed by DNA gel electrophoresis. A representative image of multiple independent experiments is shown.

We next considered repair by SSA. The reporter contains two regions of homology, i.e. the 5’ coding regions of the BFP and GFP genes, that have the potential to anneal together during SSA repair, which would result in removal of the about 3 kb intervening sequence separating the regions including the pgk-Puro gene (Fig. 3C). The annealed product would generate cells that are both BFP^-^ and GFP^-^, since the resulting BFP/GFP gene hybrid contains the C-terminally truncated region from the GFP gene (Fig. 3C). Repair by SSA would also remove the binding site for one of the primers used to PCR-amplify the Cas9 cut site for TIDER analysis (Fig. 3C, blue primer). Thus, these SSA repair products, if present, would not have been included in the TIDER analysis, resulting in a gross underestimation of the total BFP mutation frequency by this technique. Following this line of reason, we next directly compared the percentage of BFP^-^ cells detected by flow cytometry to the frequency of mutant BFP PCR products detected by TIDER analysis in both control and NU7441-treated cells. As shown in Fig. 3D, in the control DMSO-treated cells, the percentage of BFP^-^ cells was nearly identical to the percentage of mutant BFP sequences detected in the entire cell population. In contrast, in the NU7441-treated cells, the percentage of BFP^-^ cells detected by flow cytometry was significantly greater than the percentage of mutant BFP sequences detected in the TIDER analysis (Fig. 3D). Furthermore, this inability to account for all of the mutagenic repair products by TIDER provides a rationale for the paradoxical ∼10% increase in the percentage of WT sequences detected by TIDER following NU7441 treatment (Fig. 3A), even though no increase in BFP^+^ cells was observed by flow cytometry. Taken together, these results do indeed suggest an underestimation of mutagenic repair by TIDER analysis, consistent with an increase in the presence of SSA repair products following DNA-PKcs inhibition.

To directly confirm repair by SSA, we designed a PCR strategy to amplify across the entire DSB-Spectrum locus (Fig. 3C, red primers). Using this primer pair, amplification of a WT locus or a locus repaired by mutagenic end-joining or HR would generate an ∼5,000 bp product, while amplification of a locus repaired by SSA would generate a distinct ∼1300 bp fragment. Indeed, upon targeting of Cas9 to DSB-Spectrum_V2, a 1300 bp PCR product was detected (Fig. 3E), and the sequence of this PCR product aligned to the predicted SSA repair product (Fig. S3A). Importantly, the abundance of this SSA repair product was markedly increased following NU7441 treatment (Fig. 3E). Hence, inhibition of DNA-PKcs reduces c-NHEJ, which is then compensated for by both an increase in HR, and SSA.

### DSB-Spectrum_V3 is a multi-pathway DSB-repair reporter that simultaneously reports mutagenic end-joining, SSA and HR

Based on the above results, we modified DSB-Spectrum_V2 to generate a new reporter capable of distinguishing between DSB repair by HR, mutagenic end-joining, or SSA (Fig. 4A). This was achieved by inserting a pgk-promoter controlled mCherry gene in place of the pgk-Puro gene in the region separating BFP and GFP in DSB-Spectrum_V2 (Fig. 4B). With this reporter, named DSB-Spectrum_V3, repair by mutagenic end-joining will result in loss of BFP but not mCherry expression, and can therefore be distinguished from SSA which will result in loss of both fluorescent proteins (Figure 4A, B). A single copy of DSB-Spectrum_V3 was integrated into the genome of HEK 293T cells (Fig. S1C), and a single clone was expanded, which was GFP^-^, BFP^+^, and mCherry^+^, consistent with the design of the reporter. Similar to what was observed for DSB-Spectrum_V2 cells, targeting of Cas9 to DSB-Spectrum_V3 resulted in the appearance of a GFP^+^ and BFP^-^ population (Fig. 4C and D, compare AAVS1sg versus BFPsg). The BFP^-^ population could be further divided into an mCherry^+^ and mCherry^-^ population, reflecting DSB-repair by mutagenic end-joining and SSA, respectively (Fig. 4C, D). In this DSB-Spectrum_V3-expressing cell line, a surprisingly large fraction of the cells lost both BFP and mCherry expression (Fig. 4C, D). To validate that the BFP^-^ mCherry^-^ population was indeed the consequence of repair by SSA, we depleted the known SSA-factor Rad52 by RNAi. This resulted in a significant reduction in the frequency of BFP^-^, mCherry^-^ cells (Fig. 4E, F), demonstrating that DSB-Spectrum_V3 reports on SSA through loss of mCherry fluorescence.

**Figure 4.**
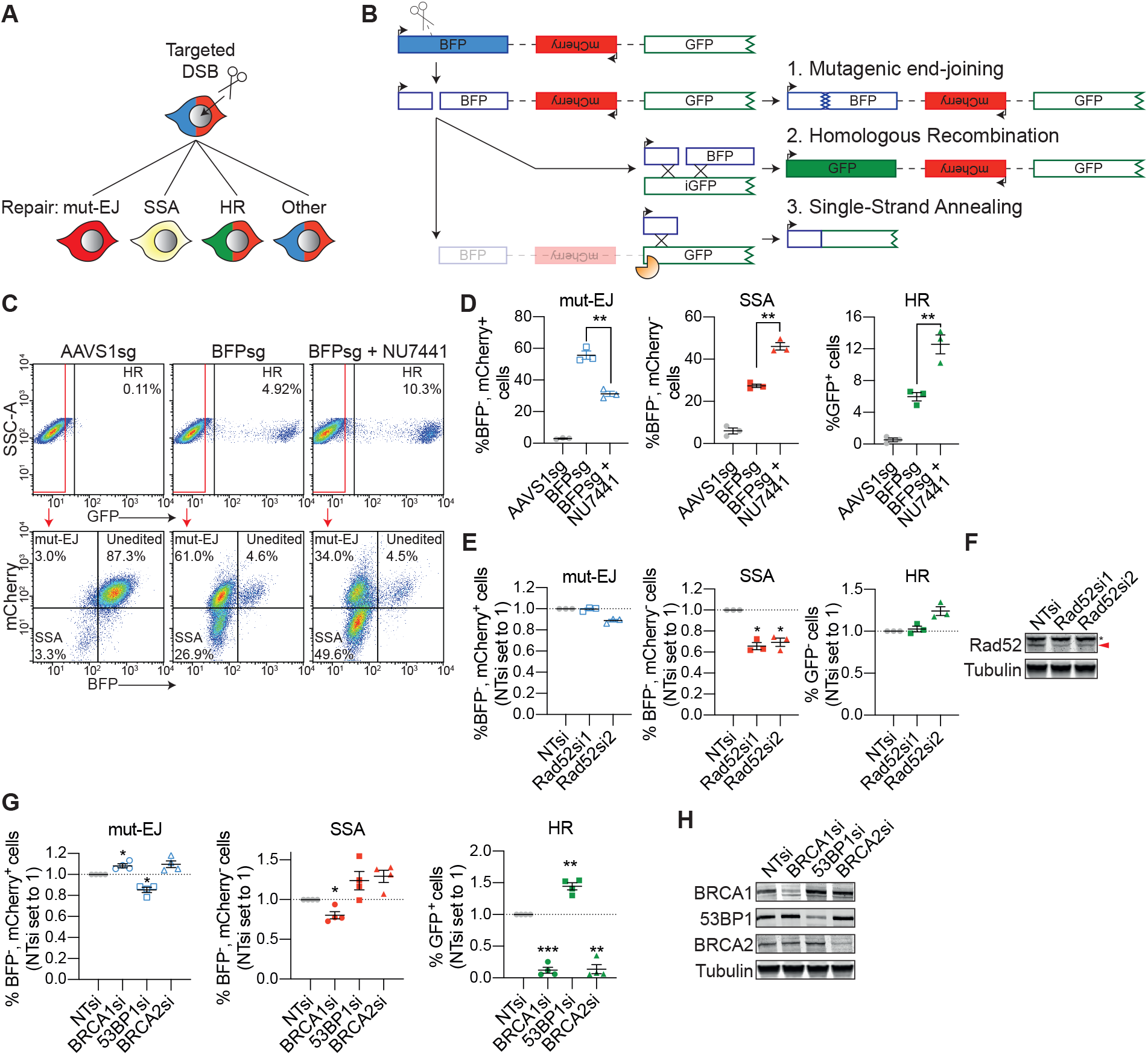
DSB-Spectrum_V3 is a reporter for mutagenic end-joining, SSA and HR. **(A)** Cartoon depicting potential outcomes of a multi-pathway reporter cell-line designed to quantify DSB-repair by mutagenic end-joining (mut-EJ), SSA and HR. **(B)** Diagramatic representation of the genomic DSB-repair reporter construct DSB-Spectrum_V3. Scissors indicate the Cas9 target site, orange pacman indicates endogenous nucleases. **(C and D)** DSB-Spectrum_V3 cells were transfected with Cas9 and an sgRNA targeting either AAVS1 or BFP, followed by treatment with NU7441 (2 µM). At 72h after Cas9 transfection cells were analyzed by flow cytometry. Panel C shows representative flow plots. Panel D shows quantification of the three repair pathways from multiple experiments (n=3; mean ± SEM; **p≤0.01). **(E)** DSB-Spectrum_V3 cells were transfected with indicated siRNAs, followed by flow cytometric analysis of mut-EJ, SSA and HR (n=3; mean ± SEM; *p≤0.05). **(F)** Western blot of lysates from cells analyzed in panel E. Tubulin is used as a loading control. Red arrow indicates Rad52 band, asterisk indicates non-specific background band. **(G)** Mut-EJ, SSA and HR was analyzed as in panel E, following siRNA-mediated knockdown of the indicated repair factors (n=5; mean ± SEM; *p≤0.05; **p≤0.01). **(H)** Western blot of lysates from cells analyzed in panel G.

Next, we used this new reporter to further examine the effects of DNA-PKcs inhibition on DSB-repair pathway choice. As shown in figures 4C, treatment with NU7441 did not affect the frequency of BFP loss, but markedly increased the frequency of mCherry^-^ cells, at the expense of mCherry^+^ cells, within the BFP^-^ population (Fig. 4C, lower panel). Quantification of multiple experiments with DSB-Spectrum_V3 cells demonstrated that DNA-PKcs inhibition increased repair by both HR and SSA while markedly inhibiting repair by mutagenic end-joining (Fig. 4D).

The unique capacity of DSB-Spectrum V3 to report on three different DSB repair pathways simultaneously provides a method to assess competition between end-joining, HR and SSA repair for the same DSB-substrate. We therefore made use of the DSB-Spectrum_V3 expressing cell line to examine crosstalk between repair pathways following perturbation of known HR and end-joining factors. First, we depleted BRCA1, 53BP1 or BRCA2 by RNAi, and quantified mutagenic-EJ, SSA and HR by flow cytometry. As expected based on its function as an end-resection inhibitor^5^, 53BP1 depletion reduced the percentage of BFP^-^, mCherry^+^ cells, but increased the percentage of both BFP^-^, mCherry^-^ cells and GFP^+^ cells, indicating suppression of mut-EJ and enhanced SSA and HR (Fig. 4G, H). Furthermore, the DSB-Spectrum_V3 reporter revealed that depletion of BRCA1 and BRCA2 strongly reduced HR and resulted in a mild promotion of mut-EJ, but had a differential effect on SSA (Fig. 4G, H). Whereas BRCA1-loss reduced SSA, depletion of BRCA2 promoted SSA (Fig. 4G, H). This finding is consistent with what has been published using a combination of individual single-pathway reporters^24^, and fits well with a model in which the end-resection function of BRCA1 promotes both HR and SSA, while the HR-specific Rad51-loading function of BRCA2 following end-resection directs repair towards HR, and away from SSA^35^. Notably, these data can also explain some seemingly puzzling results that were obtained with HEK 293T DSB-Spectrum_V2 cells. While knock-down of the end-joining factor 53BP1 resulted in suppression of mutagenic repair (fewer BFP^-^ cells) and enhancement of HR (more GFP^+^ cells; Fig. S3B, C), expected, knock-down of BRCA1 by siRNA markedly suppressed HR but surprisingly reduced, rather than enhanced, mutagenic repair, as evidenced by the reduction in both GFP^+^ and BFP^-^ cells, respectively (Fig. S3B, C). The reduced BFP-loss that we observed after knockdown of BRCA1 in the DSB-Spectrum_V2 cells can be reconciled as potentially arising from a reduction in mutagenic repair by SSA when BRCA1 is not present.

Taken together, these data validate the utility of DSB-Spectrum_V3 as a multi-pathway reporter that simultaneous quantifies repair by mutagenic end-joining, SSA and HR, with the ability to reveal both anticipated and novel DSB-repair pathway interactions.

### DSB-repair by SSA is frequent in multiple reporter contexts, genomic contexts and cell-lines

The high frequency of DSB-repair by SSA in DSB-Spectrum_V3 cells (∼27%, Fig. 4D) suggests that it can be a dominant pathway of repair, even in the presence of functional c-NHEJ and HR. To validate that this high frequency of SSA was not an artifact of the specific clonal DSB-Spectrum_V3 cell-line used, we repeated the experiments with a second monoclonal HEK 293T cell-line. A similar frequency of DSB-repair by SSA was observed in this second DSB-Spectrum_V3 clone, which further increased upon treatment with NU7441, demonstrating that the observed phenotypes are not clone-specific (Fig. S4A).

To validate the generality of the high frequency of SSA, we further quantified DSB-repair by SSA in DSB-Spectrum_V2 cells. In this reporter the homologous regions are separated by a different sequence than in DSB-Spectrum_V3, but more importantly, for this clonal cell-line we had confirmed that the reporter construct was integrated in a different genomic location than in the DSB-Spectrum_V3 clone used in figure 4 (Fig. S1C). In DSB-Spectrum_V2, SSA-repair of the DSB in the BFP gene will result in loss of the pgk-Puro cassette (Fig. 3B). SSA can therefore be quantified by measuring the number of cells that have lost puromycin resistance after generating a DSB in the reporter. We transfected HEK 293T DSB-Spectrum_V2 cells with either a control (AAVS1sg) of BFP-targeting sgRNA, and allowed gene editing to occur for 72h. Next, transfected cells were sorted by FACS, and either analyzed by flow cytometry to determine the frequency of mutagenic repair and HR, or plated to determine clonogenic survival in the absence or presence of puromycin. Using this experimental setup, we observed that ∼70% of the cells lost BFP expression and ∼4.5% underwent repair by HR (Fig. 5A). Moreover, loss of puromycin resistance was specifically observed in the BFPsg-transfected cells. On average, 22.3% of the cells lost puromycin resistance upon targeting of Cas9 to DSB-Spectrum_V2, suggesting that this fraction of cells lost the pgk-Puro cassette due to DSB-repair by SSA (Fig. 5A). Notably, this frequency of SSA is very similar to the frequency of SSA detected by mCherry loss in DSB-Spectrum_V3 cells (Fig. 4D).

**Figure 5.**
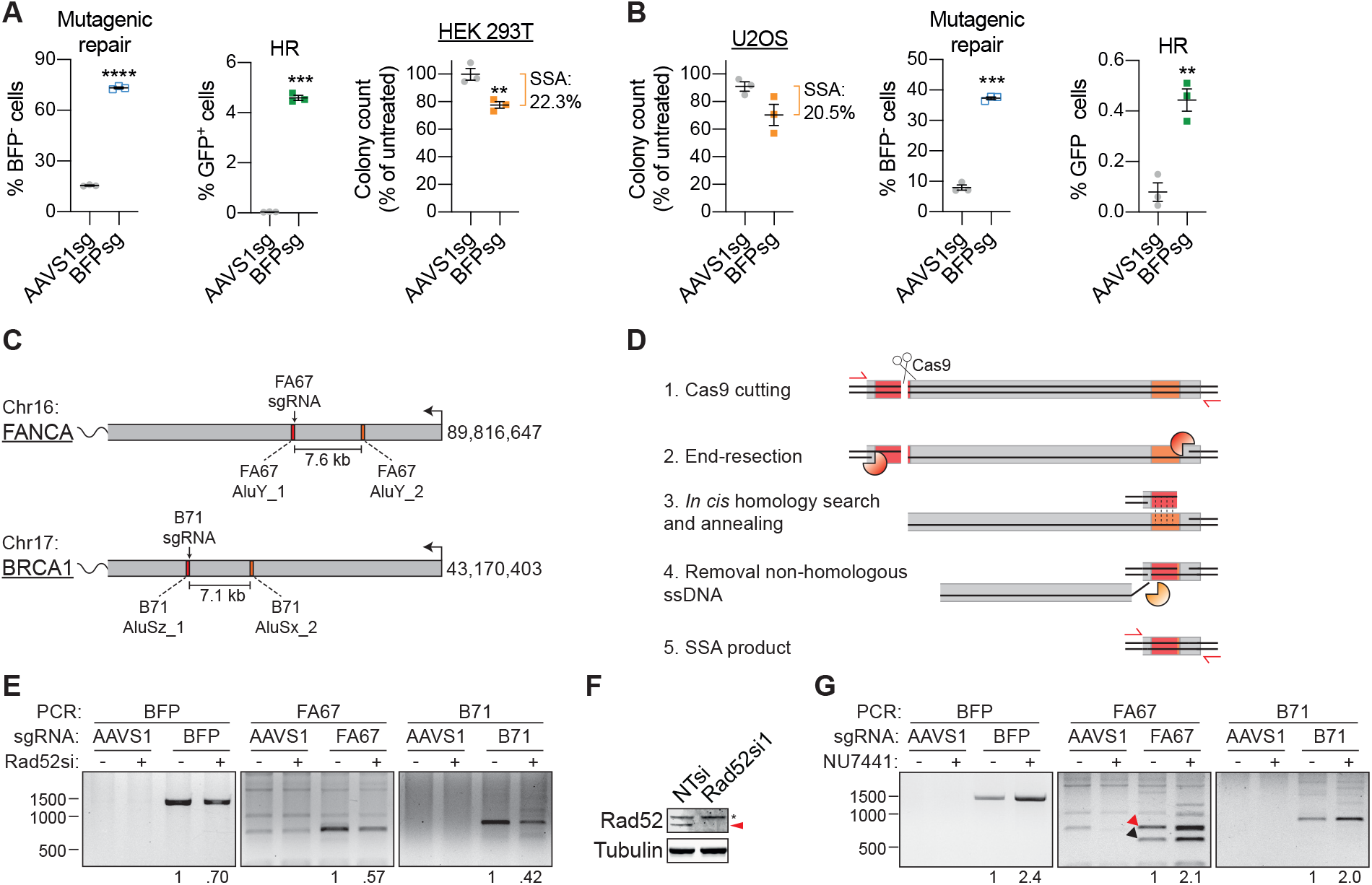
DSBs in endogenous genomic loci can be repaired by SSA between homologous Alu elements. **(A)** HEK 293T DSB-Spectrum_V2 cells were transfected with Cas9 cDNA and an sgRNA targeting either a control locus (AAVS1sg) or DSB-Spectrum_V2 (BFPsg). At 72h post-transfection, cells were harvested and either analyzed by flow cytometry to quantify mutagenic repair and HR, or replated for a clonogenic survival assay in presence or absence of puromycin, to measure SSA based on loss of the puromycin resistance gene. The number of colonies in the puromycin-treated plates were normalized to the number of colonies observed in control plates lacking puromycin (n=3; mean ± SEM; **p≤0.01; ***p≤0.001; ****p≤0.0001). **(B)** As in panel A, but for U2OS DSB-Spectrum_V2 cells (n=3; mean ± SEM; **p≤0.01; ***p≤0.001). **(C)** Schematic representation of the first part of the *FANCA* and *BRCA1* genes, which are both oriented on the minus strand of the chromosome. Indicated are the chromosomal location of the gene (according to human genome assembly hg38), the homologous Alu elements (red and orange boxes) and the sgRNA target sites. **(D)** Schematic representation of repair of the Cas9-induced DSB at the FA67 or B71 target site by SSA between the indicated Alu-elements. Indicated are the primers (in red) to amplify the SSA repair product, and the nucleases that perform end-resection (orange pacman) or ssDNA flap removal after annealing (yellow pacman). **(E)** HEK 293T DSB-Spectrum_V2 cells were transfected with a non-targeting control siRNA (in “-” lanes) or Rad52-targeting siRNA, then re-transfected with the indicated Cas9-sgRNA constructs. Genomic DNA was isolated 48h later and SSA DSB-repair products were PCR-amplified. Expected size of the respective BFP, FA67 and B71 SSA deletion products is 1249 bp, 767 bp and 906 bp. Representative pictures from two independent experiments are shown, numbers at the bottom show quantification of SSA band intensities of the presented experiment. **(F)** Western blot of lysates from panel E. Red arrow indicates Rad52 band, asterisk indicates non-specific background band. **(G)** As in panel E, but excluding siRNA transfection, and including treatment with NU7441 (2 μM). In the FA67 panel, the red arrow indicates the SSA product between the AluY_1 and AluY_2 elements, while the black arrow indicates the SSA product between the AluY_1 and a more downstream AluSx element. Representative pictures from two (B71) or more (BFP and FA67) independent experiments are shown.

Finally, to test whether SSA is also a frequently employed pathway in other cell-lines than HEK 293T, we performed the same assay in DSB-Spectrum_V2 U2OS cells. This revealed that around 20% of cells lose puromycin resistance upon targeting of Cas9 to the reporter (Fig. 5B). When compared to the 40% of total cells that had lost BFP expression and the ∼0.4% of cells that underwent repair by HR (Fig.5B), these data indicate that, also in U2OS cells, a substantial fraction of DSBs are repaired by SSA. Taken together, these results demonstrate that SSA-mediated repair of the DSB in DSB-Spectrum can be frequent in a variety of genomic contexts, and in multiple cell-lines.

### DSBs at endogenous genomic loci can be repaired by SSA through annealing of homologous Alu elements

The DSB-Spectrum reporters are potentially prone to SSA-mediated repair due to the highly homologous BFP and GFP sequences within 3 kb of the DSB (Fig. 3B). We therefore investigated whether SSA is also employed to repair DSBs in endogenous genomic loci. It has been suggested that Alu elements can function as homologous regions in the human genome that anneal during DSB-repair by SSA^9^, because these 200-300 bp elements are abundant^36^, and can share high levels of sequence homology. To test this hypothesis, we designed an sgRNA to target Cas9 to a site within an Alu element in the endogenous *FANCA* gene. This element (AluY_1) shares 86% homology with a second Alu element (AluY_2) located 7.6 kb downstream from the first (FA67 sgRNA; Fig. 5C, S4B). Upon Cas9-induced DSB generation, long range end-resection could potentially expose both homologous AluY elements, which in turn could anneal during SSA, followed by removal of the 7.6 kb region between the AluY elements (Fig. 5D). The presence of such an SSA-induced deletion product, if created, could then be detected by PCR (Fig. 5D). To test a second independent site in a different gene, we designed a different sgRNA to target Cas9 to an AluSz element in the endogenous *BRCA1* gene, that is highly homologous to an AluSx element located 7.1 kb downstream (B71 sgRNA; Fig. 5C, S4C).

We transfected HEK 293T DSB-Spectrum_V2 cells with Cas9 cDNA and an sgRNA, to target a control locus (AAVS1), the site in the *FANCA* gene (FA67), the site in the *BRCA1* gene (B71) or DSB-Spectrum (BFP). We next isolated genomic DNA, and performed PCR to detect the deletion product generated by SSA-mediated repair of the target loci (Fig. 5D). The size of the PCR product was predicted to be 767 bp, 906 bp and 1249 bp for the FA67, B71 and BFP SSA deletion products, respectively. For all three loci, we could readily detect a specific PCR product of the expected size, only when cells were transfected with the FA67, B71, or BFP sgRNAs, but not when they were transfected with the AAVS1-targeting control sgRNA (Fig. 5E). Sequencing of these PCR products confirmed that they corresponded to the predicted SSA DSB-repair products (Fig.S5A, B). Moreover, upon knockdown of the SSA-factor Rad52, the yield of these PCR products was reduced, confirming that they were the consequence of SSA-mediated repair of the Cas9-induced DSB (Fig. 5E, F). These results show that Alu elements, of which there are over a million interspersed throughout the human genome^36^, can drive DSB-repair by the efficient but highly mutagenic SSA pathway.

Our results with DSB-Spectrum reporters indicated that inhibition of DNA-PKcs strongly promotes SSA (Fig. 3C, 4D). To test whether this is also the case for SSA-mediated repair of endogenous genomic loci, we performed FA67 and B71 editing experiments as described above, but now included treatment with NU7441 for the duration of the gene editing. As shown in figure 5G, the SSA DSB-repair product was specifically detected upon DSB-generation in the FA67 and B71 target loci, as well as in the BFP positive control locus. Moreover, for all three loci these products were over two-fold more abundant when DNA-PKcs was inhibited by NU7441 treatment (Fig. 5G). Interestingly, in this experiment, for the FA67 locus, a second, smaller PCR product was observed specifically in the FA67sg-transfected cells, and this smaller-sized product was also increased in abundance upon NU7441 treatment (Fig. 5G). Sequence analysis of this PCR-product demonstrated it to be the consequence of SSA between the AluY_1 element and an AluSx element directly downstream of the AluY_2 element, generating a larger deletion and thus a smaller SSA-repair PCR product (Fig. S4D, E, S5A). Together, these data demonstrate that DSBs in endogenous genomic loci can be repaired by SSA between homologous Alu elements, and that this mutagenic repair is strongly stimulated by inhibition of DNA-PKcs kinase activity.

### Inhibition of long-range end-resection promotes HR and reduces SSA

The use of DNA-PKcs kinase inhibitors has been suggested as an HR-promoting strategy in CRISPR-mediated genome editing experiments^37,38^. Our data indicate that DSB-repair by SSA is strongly promoted by these drugs (Fig. 5C-G). Since SSA can result in loss of kilobases of DNA^12^, unbalanced levels of SSA could have profound consequences on genomic integrity. Thus, when applying DNA-PKcs inhibition in the context of genome editing, in a research setting but particularly in the clinic, it will be essential to prevent the SSA-promoting effects of this treatment. We therefore investigated DSB-repair pathway choice between SSA and HR using DSB-Spectrum_V3. Although repair by both SSA and HR requires extensive end-resection, data from Ochs *et al*. indicated that hyper-resection may promote SSA over HR^39^. We therefore investigated whether there was a differential requirement for specific end-resection factors for SSA compared to HR. The short-range end-resection factors Mre11 or CtIP, or the long-range end-resection factors Exo1 or DNA2 were depleted by RNAi, and the modes of DSB-repair in DSB-Spectrum_V3 cells were analyzed by flow cytometry. As shown in Figure 6A and B, loss of Mre11 and CtIP resulted in a reduction of DSB-repair by both SSA as well as HR, consistent with a requirement for end-resection for both pathways (Fig. 6A, B). A similar reduction in SSA was observed upon knockdown of Exo1 and DNA2, but surprisingly, loss of these factors promoted, rather than reduced, HR (Fig. 6A-C). Thus, the efficiency of DSB repair by SSA, but not HR, is dependent on factors involved in long-range end-resection, and these factors can even inhibit HR.

**Figure 6.**
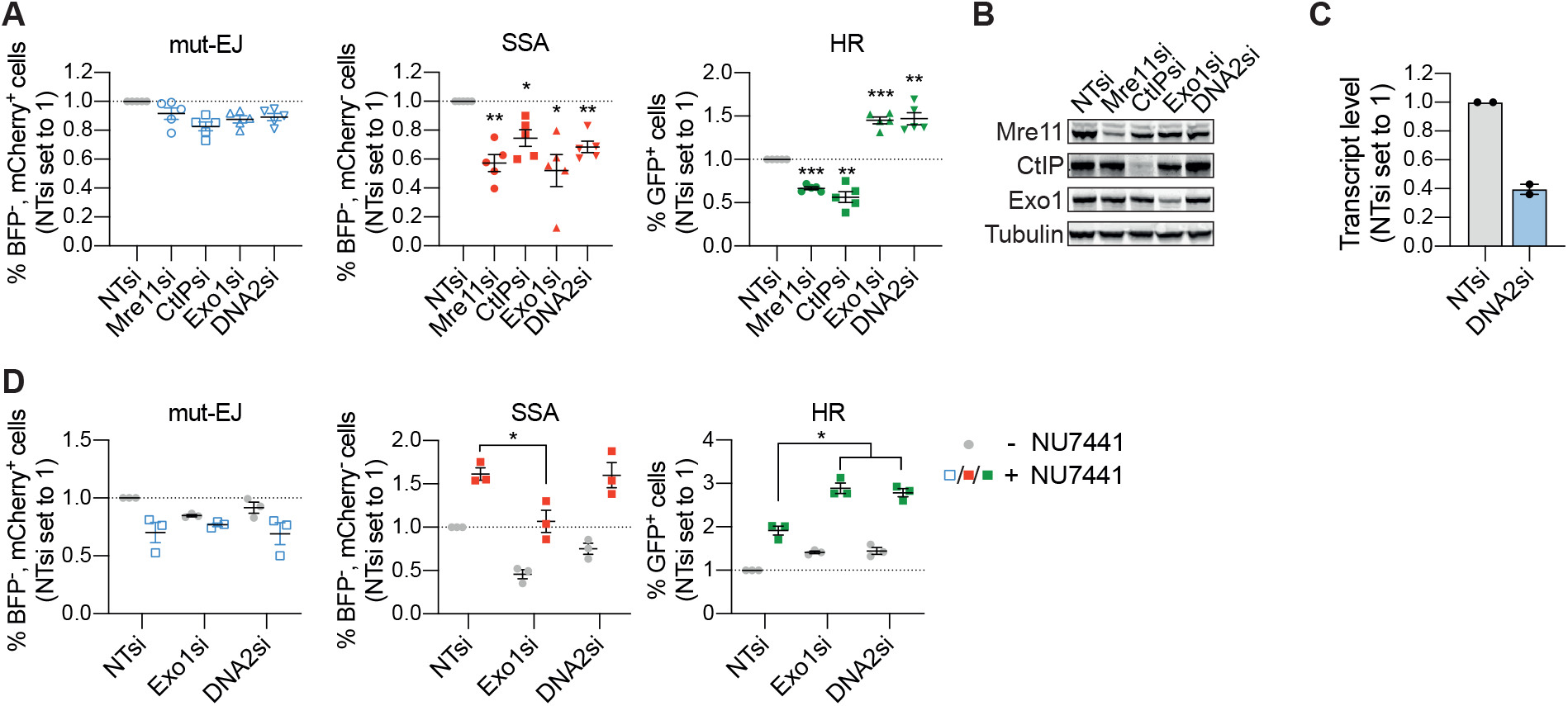
The long-range end-resection factors Exo1 and DNA2 are required for SSA, but inhibit HR in DSB-Spectrum_V3 cells. **(A)** DSB-Spectrum_V3 cells were transfected with indicated siRNAs, followed by transfection with Cas9 and an sgRNA targeting a control locus or BFP. At 72h after Cas9 transfection cells were analyzed by flow cytometry, and repair pathways quantified as figure 1E (n=5; mean ± SEM; *p≤0.05; **p≤0.01; ***p≤0.001). **(B)** Western blot of lysates from cells analyzed in panel A. Tubulin is used as a loading control. **(C)** Quantification of DNA2 knock-down in cells used in panel A. RNA was isolated, followed by reverse transcription and quantitative PCR. The qPCR reaction was performed using two independent primer pairs, each symbol represents the average of a technical duplicate with one of the two primer pairs. **(D)** As in panel A, including treatment with NU7441 (2 µM) at 16h after Cas9 transfection (n=3; mean ± SEM; *p≤0.05).

This finding prompted us to test, using DSB-Spectrum_V3 cells, whether knockdown of either Exo1 or DNA2 would prevent the SSA-promoting effect of DNA-PKcs inhibition. As shown in Fig. 6D, in the absence of DNA-PKcs inhibition (grey symbols), knockdown of Exo1 or DNA2 reduced SSA and promoted HR. Notably, the SSA-reducing effect of DNA2 knockdown was smaller than that seen with Exo1 knockdown (Fig. 6A, D). Upon treatment with NU7441, mutagenic end-joining (BFP^-^/mCherry^+^ cells) was reduced, as expected, and this effect was essentially unchanged by knock-down of either Exo1 or DNA2 (Fig. 6D). Importantly, however, the increase in HR following DNA-PKcs inhibition was enhanced by ∼50% when Exo1 or DNA2 was depleted (Fig. 6D). Furthermore, in these experiments, Exo1 depletion, but not DNA2 depletion, blunted the increase in SSA that was observed following NU7441 treatment, resulting in a frequency of SSA repair after DNA-PKcs inhibition that was similar to the frequency in untreated cells (Fig. 6D). We do not understand the cause of the differential effects of Exo1 and DNA2 on SSA in the presence of NU7441, but the data suggest that the relationship between end-resection and DBS-repair pathway choice is likely complex. Nevertheless, our findings indicate that in DSB-Spectrum_V3, DSB-repair pathway choice can be efficiently channeled towards HR without promotion of mutagenic SSA, by concomitantly inhibiting both c-NHEJ and long-range end-resection by Exo1.

## Discussion

In this manuscript we describe and validate three distinct reporter systems for DNA DSB-repair, DSB-Spectrum_V1, V2, and V3, capable of flow cytometric quantification of repair by error-free c-NHEJ versus HR in case of V1, mutagenic repair versus HR in case of V2, or mutagenic end-joining, versus HR versus SSA in case of V3. These DSB-Spectrum reporters build on a large history of DSB-repair reporter constructs, and their design is based on commonly used single-pathway reporter systems such as DR-GFP and variations there-of^23–26^. These single-pathway reporters, designed by others, have proven extremely valuable in DSB-repair research^40^. The DSB-Spectrum reporters that we describe here have the added advantage that they can simultaneously quantify the frequency of DSB-repair by multiple different pathways within a single population. This facilitates the analysis of pathway crosstalk, and reveals DSB-repair pathway interdependence and compensation as they compete for the same DSB-substrate.

A number of fluorescence-based two-pathway reporters have been described^28,41–45^. Most of these published two-pathway reporters require the addition of an exogenous repair template to study DSB-repair by HR, rendering them technically more challenging to use than DSB-Spectrum reporters^28,42,44,45^. Moreover, DSB_Spectrum_V3 has the unique capacity to report on three DSB repair pathways, which is only shared with one other reporter system, the SSA-TLR reporter developed by Certo, Scharenberg, and colleagues^27^. That reporter can quantify mutagenic end-joining, SSA and HR, like DSB-Spectrum_V3. However, the frequency of detected end-joining events by the SSA-TLR reporter is very low compared to DSB-Spectrum, because the SSA-TLR can only detect these events if the resulting InDels shift the reading frame by two bp^27^. Finally, in case of the SSA-TLR, the detected frequency of HR events is strongly dependent on the amount of repair template co-delivered with the nuclease, and in the majority of assays presented by the authors the HR-frequency is lower than 0.5%^27^. Thus, we believe that the technical simplicity, high efficiency, and repair pathway inclusivity make DSB-Spectrum reporters ideal tools to study DSB-repair on the network level.

Our finding that in both HEK 293T and in U2OS cells, the frequency of detected DSB repair by SSA was substantially higher (20-25%) than the frequency of detected repair by HR (0.5-5%) is surprising but entirely consistent with the high frequencies of DSB-repair by SSA that have been reported using other reporter systems^11,27,41,46^. Two of these studies allowed direct comparison between SSA and other repair pathways. Jasin and colleagues analyzed DSB-repair products following endonuclease cleavage of a genomic reporter construct in CHO cells, and found that repair products consistent with SSA were twice as frequent as those generated by HR^46^. Huertas and colleagues used a fluorescent SSA versus c-NHEJ reporter and found comparable levels of repair by both of these two pathways^41^. In the vast majority of reporter systems of DSB repair, including DSB-Spectrum, the presence of highly homologous sequences adjacent to the DSB generates an optimal sequence context for SSA, which may predispose to repair by this pathway. Nevertheless, these data, taken together, demonstrate that repair by SSA can be very common in the right sequence context, despite its highly mutagenic consequences.

This SSA-prone sequence context for DSB repair might not be rare in the human genome, which contains a large number of repetitive sequences^47^, including over a million Alu elements^36^. Interestingly, a study of familial mutations in the *FANCA* gene, which causes the disease Fanconi Anaemia (FA), found that 75% of a total of 88 pathogenic deletions were between two Alu elements^48^. Similarly, deletions between Alu elements have also been reported in *BRCA1* mutant carriers^49,50^, who have an increased risk of developing breast and ovarian cancer^51^. Here we show that a Cas9-induced DSB within the endogenous genomic loci of the *FANCA* or *BRCA1* genes can be repaired by SSA through annealing of adjacent homologous Alu elements (Fig. 5C-G). The SSA-induced deletion between the AluY_1 and AluY_2 elements in the *FANCA* gene was nearly identical to the pathogenic deletion identified in one of the Fanconi Anaemia families (ID 67)^48^. This was also the case for the SSA-induced deletion between the AluSx and AluSz elements in the *BRCA1* gene, which strongly resembled the deletion identified in a family with a germline *BRCA1* mutant allele^50^. The exact molecular mechanism responsible for the Alu-Alu deletions seen in the familial *FANCA* or *BRCA1* mutants described above is not known. However, our findings indicate that SSA-mediated repair of a DSB in the vicinity would have the potential to generate such deletion products.

Our studies also demonstrated that DSB-repair by SSA is strongly promoted by inhibition of DNA-PKcs kinase activity using NU7441. A slight increase in SSA upon treatment with NU7026, another DNA-PKcs kinase inhibitor, has previously been reported, demonstrating that this phenotype is not dependent on the specific small molecule inhibitor used^52^. This SSA-promoting effect is important because inhibition of DNA-PKcs has been used to increase the HR to c-NHEJ ratio in the context of CRISPR-mediated genome editing^8,37,38^. A concurrent upregulation of SSA has not been addressed by these studies and large deletion products would, in most cases, have been missed by next generation sequencing of a small region surrounding the DSB junction^53^. Our data therefore argue that quantification of repair by SSA should be included in gene editing and DSB-repair manipulation studies, since an increase in the highly mutagenic repair by SSA can have deleterious consequences for genomic stability. Furthermore, several studies in cell-lines, and in human embryos, have demonstrated that a single Cas9-induced DSB can result in kilobase-sized genomic deletions^53–55^. It would be interesting to examine whether SSA is responsible for these large deletions.

To provide a potential solution for the high levels of SSA following inhibition of end-joining, we used DSB-Spectrum_V3 cells to study DSB-repair pathway crosstalk between end-joining, HR and SSA. We found that optimal SSA requires the long-range end-resection factors Exo1 and DNA2, as anticipated on basis of the 3 kb distance between the homologous regions in DSB-Spectrum_V3. This is substantially larger than the short-range resection-span of 200-300 nucleotides typically generated by Mre11 and CtIP^56^. To our surprise, our studies with DSB-Spectrum_V3 showed that loss of Exo1 and DNA2 promoted HR, which contradicts the conventional view of end-resection and pathway choice, suggesting that Exo1 and DNA2 are positive regulators of HR^57^. However, our data are in good agreement with published reports showing an increase in HR upon DNA2 knockdown, and a decrease in HR upon hyperactivation of Exo1^58,59^. Furthermore, Chen *et al*. showed that reconstitution of Exo1 knock-out cells with Exo1 cDNA reduced HR, especially between repeats with some divergence in sequence^60^. Lastly, depletion of 53BP1 induces hyper-resection, which correlates with a DSB-repair switch from Rad51-dependent HR to Rad52-dependent SSA^39^. Hence, our data, together with these previously published results, fits best with a model in which long-range end-resection is not absolutely required for HR and might, in some cases, direct DSB-repair towards SSA instead.

Taken together, our studies using the novel DSB-repair reporters DSB-Spectrum_V1 to V3 demonstrate that inhibition of DNA-PKcs promotes SSA as well as HR, and that loss of Exo1 or DNA2 can channel DSB-repair away from SSA towards HR. Multi-pathway analysis using DSB-Spectrum facilitates the generation of an inclusive model of DSB-repair pathway choice (Fig. 7), and provides a comprehensive understanding of DSB-repair outcomes following perturbation of the DSB signaling and repair network.

**Figure 7.**
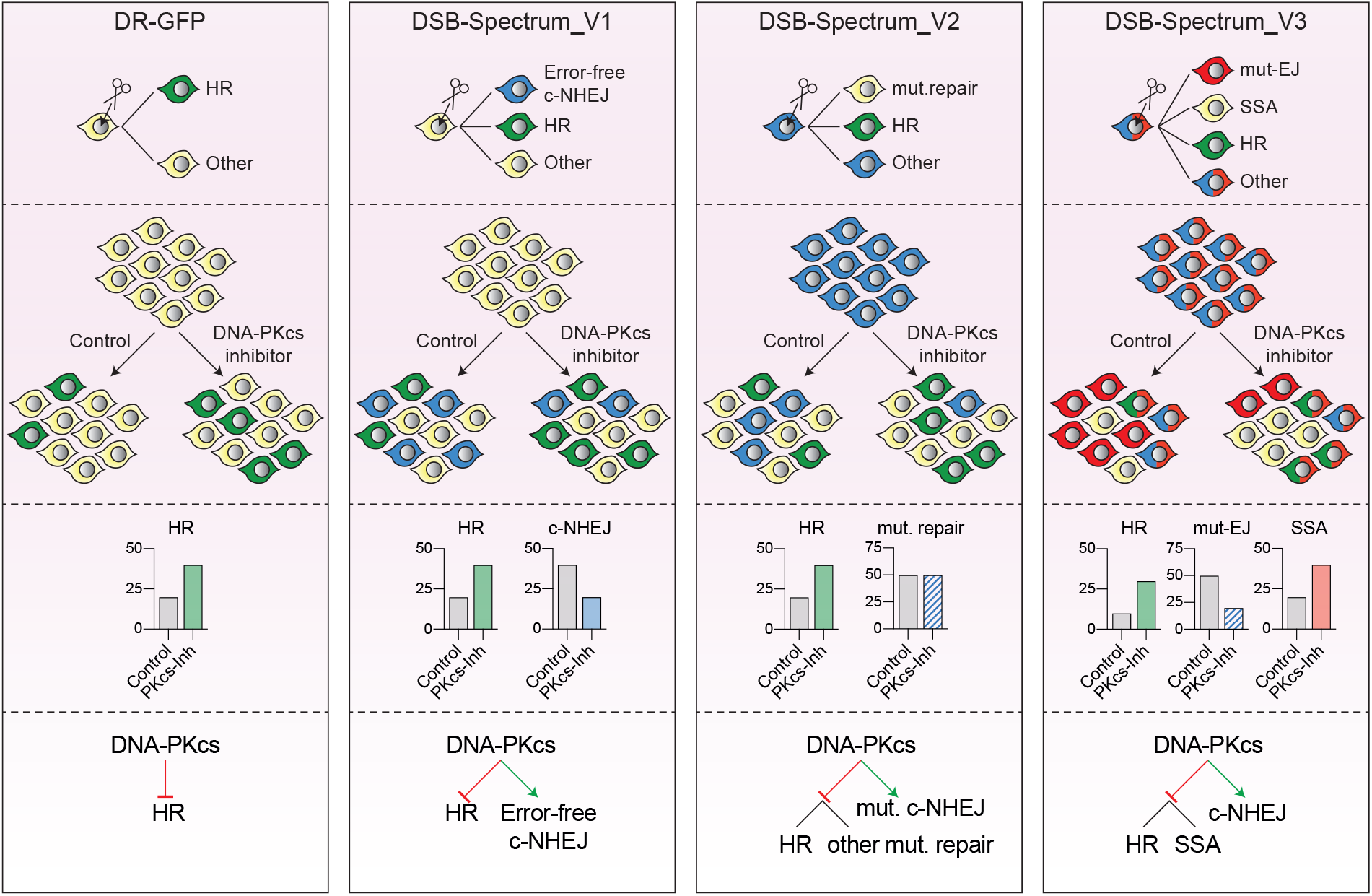
Multi-pathway quantification of DSB-repair using DSB-Spectrum allows for comprehensive analysis of repair outcomes in absence and presence of DSB-repair manipulation strategies. Summary of the DSB-Spectrum variants described in this manuscript. The frequently used DR-GFP reporter is shown for comparison purposes. Each panel shows, from top to bottom (1) the fluorescence changes caused by repair of the site-specific DSB in the depicted reporter construct, (2) a cartoon example of a population after reporter activation, in presence or absence of DNA-PKcs inibition as an example of a DSB-repair manipulation strategy, (3) quantification of the results in panel 2, (4) the model resulting from the data obtained with the reporter construct.

## Materials and Methods

### Cloning

All cloning was done using standard PCR-protocols, restriction/ligation protocols, or fragment assembly using the NEBuilder HiFi DNA Assembly Master Mix (New England Biolabs) according to manufacturer’s instructions. DNA plasmids were purified using Plasmid Mini/Midi or Maxi kits (Qiagen) according to manufacturer’s instructions. The DSB-Spectrum_V2 construct was generated in multiple steps. First, the gene encoding for eBFP1.2 was generated by multiple rounds of site-directed mutagenesis of pEGFP-C1 (Clontech). Next, an XhoI restriction site was generated 5’ of the GFP-gene in pDRGFP (Addgene plasmid #26475)^23^ and the GFP-gene was replaced by the eBFP1.2 gene using the XhoI/NotI restriction sites. Finally, the region from the start of eBFP1.2 until the end of iGFP was PCR-amplified from the modified pDRGFP and ligated into XbaI/MluI digested pLVX-IRES-Hyg (Clontech; this procedure removes the IRES-Hygro sequence). To generate DSB-Spectrum_V1, the spacer sequence was introduced into eBFP1.2 by PCR, and wild-type eBFP1.2 in DSB-Spectrum_V2 was replaced by spacer-separated eBFP1.2 using fragment assembly. To generate DSB-Spectrum_V3, the puromycin resistance gene in DSB-Spectrum_V2 was replaced by an mCherry gene using fragment assembly. All DSB-Spectrum plasmid propagation was done in recombination-deficient bacteria (Stbl3; ThermoFisher Scientific).

To generate the Cas9/sgRNA plasmids, the puromycin resistance gene in pSpCas9(BB)-2A-Puro (PX459; Addgene plasmid #62988)^61^ was replaced by either mCherry or iRFP670 using PCR and fragment assembly. A PCR-strategy was designed to destroy the BpiI sites in mCherry and iRFP670. Cloning of sgRNAs was done using the BpiI-sites in pX459 as described in Ran et al^61^. The sgRNA target sequences were as follows: AAVS1: 5’-GGCAAAATTCCCTCAGTTTA-3’, BFP sgRNA V2: 5’-ACGGGGTCCAGTGTTTTGCC-3’, BFP sgRNA 1: 5’-ACCACCCTCTCCCACGGCTA-3’, BFP sgRNA2: 5’-GGGCAAAACACTGGACCAAT-3’, FANCA FA67 sgRNA: 5’-GAAAGAGCCAGACTCCGTCT-3’, BRCA1 B71 sgRNA: 5’-GGGAGGCAGACGTTGCGGAG-3’. The AAVS1 sgRNA targets intron 1 of regulatory subunit 12C gene of protein phosphatase 1 (PPP1R12C), located on human chromosome 19. The targeting vector for DSB-Spectrum_V1, containing two sgRNAs, was generated by introducing each individual sgRNAs into a separate pX459-mCherry vector. Subsequently, the U6-sgRNA region of sgRNA 1 was put into pX459-mCherry with sgRNA 2 using XbaI/KpnI. All constructs were sequence verified by sanger sequencing.

### Generation of DSB-Spectrum cell-lines

Cell-lines were grown at 37°C in a humidified incubator supplied with 5% CO2, in DMEM supplemented with 10% FBS. To produce lentivirus, 3*10^6^ HEK 293T cells were plated in 10 cm plates and transiently transfected with pLVX-DSB-Spectrum and packaging vectors (pCMV-VSVg and pCMV-ΔR8.2) using the CalPhos mammalian transfection kit (Clontech). Virus-containing medium was filtered through a 0.45 μM filter, supplemented with PolyBrene (8 μg/ml) and transferred to a 10 cm plate with HEK 293T or U-2 OS target cells at 70% confluency. Due to the large size of pLVX-DSB-Spectrum the viral titer was low, so this strategy resulted in infection with low MOI (<0.2). In the case of DSB-Spectrum V1 and V2, infected cells were cultured on selection medium containing puromycin (1 μg/ml; Invivogen). After selection, BFP^-^/GFP^-^ (V1), BFP^+^/GFP^-^ (V2) or BFP^+^/mCherry^+^/GFP^-^ (V3) cells were sorted and single-cell plated in 96-well plates by FACS, followed by expansion of single-cell clones. To test the expanded clones for efficiency of Cas9 cleavage, they were transfected with Cas9 and sgRNA(s) targeting DSB-Spectrum, and 48-96h later fluorescent populations were quantified by flow cytometry. Clones for which the anticipated fluorescent populations could be detected were selected and propagated as DSB-Spectrum cell-lines.

### DSB-Spectrum assays

DSB-Spectrum cells were plated and transfected the next day with pX459-Cas9-sgRNA-mCherry or pX459-Cas9-sgRNA-iRFP constructs containing either an AAVS1 sgRNA or sgRNA(s) targeting DSB-Spectrum. Transfection was done using Lipofectamine 2000 (Invitrogen) according to manufacturer’s protocol. At 48-96h after transfection, cells were trypsinized and analyzed by flow cytometry. For experiments involving RNAi, cells were plated and transfected with siRNA the next day, followed by replacement of transfection medium and a second siRNA transection 24h after the first siRNA transfection. Transfection was performed using Lipofectamine RNAiMAX (Invitrogen) according to manufacturer’s protocol. Cells were re-plated at 6-8h after the second siRNA transfection, and transfected with pX459-Cas9-sgRNA constructs 24h after replating. A slightly different protocol was used for the experiments with BRCA1 and 53BP1 knockdown in HEK 293T DSB-Spectrum_V2 cells (Fig. S3B). In this case, only one round of siRNA transfection was done, and pX459-Cas9-sgRNA constructs were transfected at 48h after siRNA transfection. All siRNAs were Ambion Silencer Select Pre-designed siRNAs (Life Technologies) with the following ID#: s11746 (Rad52si1), s11747 (Rad52si2), s224682 (BRCA1si), s14313 (53BP1si), s773 (DNA-PKcssi), s2085 (BRCA2si), s4173 (DNA2si), s11849 (CtIPsi), s17502 (Exo1si), s8959 (Mre11si), s14952 (Ku80si), s14949 (XRCC4si), s8179 (Lig4si) and s21059 (PolQsi). Treatment with NU7441 (2 μM; SelleckChem) was done either directly before Cas9-sgRNA transfection or 16h afterwards. Following flow cytometry, the different DSB-repair frequencies were quantified using FlowJo software (BD Biosciences). Gating on forward and side-scatter was applied to select the live, single-cell, population, followed by gating on either mCherry or iRFP to select transfected cells. On this population care full gating was applied to quantify the frequencies of BFP^+^ and GFP^+^ (DSB-Spectrum V1), BFP^-^ and GFP^+^ (DSB-Spectrum V2), or BFP^-^/mCherry^-^, BFP^-^/mCherry^+^ and GFP^+^ (DSB-Spectrum V3) cells. These frequencies are plotted directly in Figures 1D, 1I, 2D, 2G, 4D and supplementary figure 4A. For all other experiments, the frequency of each fluorescent sub-population in the AAVS1sg- transfected cells was subtracted from the frequency of that same population in the BFPsg-transfected cells, and the resulting background-corrected frequencies were normalized to the NTsi-cells.

### TIDE(R)-analysis and SSA-product PCR

For TIDE(R) analysis and PCR-mediated detection of the SSA repair product, a DSB-Spectrum assay was performed, and genomic DNA isolated using the DNeasy Blood and Tissue kit (Qiagen) according to manufacturer’s instruction. The DSB-Spectrum target region was then PCR-amplified using the following primers: TIDE Forward 5’-CGTAACAACTCCGCCCCATT-3’ and TIDE Reverse 5’-GGGTGTTCTGCTGGTAGTGG-3’, or SSA Forward 5’-CGGTTTGACTCACGGGGATTTC-3’ and SSA Reverse 5’-CGGGCCACAACTCCTCATAAAG-3’. the TIDE PCR product was submitted for Sanger sequencing, either directly from the PCR-reaction mix, or after PCR purification using the QiaQuick PCR purification kit (Qiagen) according to manufacturer’s instruction. Sequence files were uploaded for TIDE(R) analysis on (tide.nki.nl). The SSA-PCR product was analyzed by agarose gel electrophoresis, and similarly sequenced by Sanger sequencing.

### Western blotting and antibodies

Samples for western blotting were harvested 72h-96h after the first round of siRNA transfection. Cells were lysed on ice in RIPA buffer (50 mM Tris-HCl pH 8.0, 1 mM EDTA, 1% Triton-X100, 0.5% Sodium Deoxycholate, 0.1% SDS, 150 mM NaCl) supplemented with cOmplete EDTA-free Protease Inhibitor Cocktail tablets (Roche), 2 mM MgCl2 and Benzonase Nuclease (100 U/ml; Merck Millipore). Lysates were cleared by centrifugation at 14,000 rpm for 15 minutes, and protein concentration was determined by a BCA assay (Pierce). Next, SDS-sample buffer (pH 6.8) was added to the lysates, followed by incubation at 95°C for 5 minutes. Subsequently, equal amounts of protein for each sample was loaded on 4-15% Criterion TGX pre-cast midi protein gel (Bio-Rad). After SDS-PAGE, proteins were transferred to nitrocellulose membranes using a standard tank electrotransfer protocol. Membranes were blocked using either 5% skim milk or Blocking buffer for fluorescent WB (Rockland) in PBS, followed by antibody staining in Blocking buffer for fluorescent WB (Rockland) diluted 1:1 in PBS with 0.1% Tween-20. After secondary antibody-staining, the membranes were imaged on an Odyssey CLx scanner (LI-COR BioSciences). Image analysis was done using ImageStudio (LI-COR BioSciences). Primary antibodies used were Rabbit*α*BRCA1 (Millipore 07-434; 1:2000), Rabbit*α*53BP1 (Novus NB100-304; 1:2000), Rabbit*α*Ku80 (Santa-Cruz sc-9034; 1:200), Mouse*α*DNA-PKcs (Abcam ab44815, 1:700), Mouse*α*XRCC4 (Signalway antibody 40455, 1:1000), Rabbit*α*Ligase-4 (Abcam ab 193353, 1:1000), Mouse*α*Tubulin (Sigma T6199, 1:3000), Mouse*α*BetaActin (Sigma, 1:10,000), Mouse*α*BRCA2 (Merck-Millipore OP95, 1:500), Mouse*α*Rad52 (Santa-Cruz sc-365341, 1:100,), Rabbit*α*Exo1 (Abcam ab95068; 1:2000), Mouse*α*CtIP (Millipore MABE1060; 1:1000), and Rabbit*α*Mre11 (Kind gift from prof. Roland Kanaar, Erasmus MC, Rotterdam, the Netherlands; 1:3000)^62^. Secondary antibodies used were Goat*α*Mouse or Goat*α*Rabbit labeled with either IRDye 680 or IRDye 800 (LI-COR, 1:10,000).

### Clonogenic survival assays

HEK 293T DSB-Spectrum_V2 or U2OS DSB-Spectrum_V2 cell lines were transfected with pX459-Cas9-sgRNA-iRFP constructs containing either the AAVS1 targeting control sgRNA of BFP-targeting sgRNA 2. Transfections were done with Lipofectamine 2000 (Invitrogen) according to manufacturer’s instructions. At 48h after transfection, iRFP-positive cells were collected using FACS and plated in 6-wells plates, either 400 cells per well for U2OS cells, or 800 cells per well for the HEK 293T cells. The HEK 293T cells were plated in Poly-L-lysine coated plates to prevent detachment during later washing and staining steps. Of note, while performing the FACS to collect iRFP-positive cells, analysis of BFP and GFP expression in this population was also performed to determine the frequency of mutagenic repair and HR. Directly after plating, cells were either left untreated or treated with puromycin (2 μg/ml; Invivogen). At 10d after plating, cells were washed with PBS and stained and fixed with staining solution containing 0.4% Crystal Violet and 20% methanol. Plates were scanned and colonies were counted using ImageJ.

### SSA between Alu elements

To study SSA between Alu elements at endogenous loci in the *FANCA* and *BRCA1* genes, experiments were performed in HEK 293T DSB-Spectrum_V2 cells essentially as described for the DSB-Spectrum assays. However, cells were transfected with FA67 and B71 sgRNAs, in addition to AAVS1 (negative control) and BFP-targeting sgRNAs (positive control). Also, no flow cytometric analysis was performed, but instead cells were harvested at 48h-72h after Cas9-sgRNA transfection, followed by isolation of genomic DNA using the DNeasy Blood and Tissue kit (Qiagen) according to manufacturer’s instruction. Next, the SSA repair product was PCR-amplified using GoTaq G2 polymerase (Promega), in 25 μl reaction volumes containing 100-250 ng genomic DNA as template. When product yield was compared between samples, the exact amount of genomic DNA template was equalized. The following primers were added to an end-concentration of 0.2 μM per primer: FA67 SSA FWD: 5’-TTACAGTCTGGGCTGCAGTG-3’, FA67 SSA REV: 5’-AAAGCCCAGAATCAGACGGG-3’, B71 SSA FWD: 5’-TCAGTGCCTGTTAAGTTGGC-3’, B71 SSA REV: 5’-CTGGCAACATCTCTTTATTGAGCA-3’, or the BFP SSA primers described above. To amplify the FA67 and B71 SSA repair products, the following cycling parameters were used: 1. 95°C 02:00 – 2. 95°C 00:15 – 3. 68°C (−1°C per cycle) 00:15 – 4. 72°C 01:30 – Goto step 2 10x – 5. 95°C 00:15 – 6. 57.4°C 00:15 – 7. 72°C 01:30 – Goto step 5 26x - 72°C 10:00 – Hold at 10°C. To amplify the SSA repair product from the BFP locus, DMSO was added to the PCR reaction mixture to an end-concentration of 5%. In addition, the following cycling parameters were used: 1. 95°C 02:00 – 2. 95°C 00:15 – 3. 68°C (−1°C per cycle) 00:15 – 4. 72°C 01:30 – Goto step 2 10x – 5. 95°C 00:15 – 6. 55°C 00:15 – 7. 72°C 01:30 – Goto step 5 20x - 72°C 10:00 – Hold at 10°C. After PCR, reaction mixtures were directly analyzed by DNA gel electrophoresis.

### Splinkerette PCR

To identify the genomic integration sites of the DSB-Spectrum reporter cassettes, a Splinkerette PCR protocol was performed adapted from Yin *et al*^29^. In short, genomic DNA was isolated using the DNeasy Blood and Tissue kit (Qiagen) according to manufacturer’s instruction. Next, 2 μg of genomic DNA was digested with TfiI (NEB) and StyI (NEB), incubating for 6h at 37°C and another 6h at 65°C. The digested DNA was purified from the restriction mix using the PCR purification kit (QiaGen) according to manufacturer’s instruction, followed by incubation with dNTPs and GoTaq G2 polymerase (Promega) at 72°C for 30 minutes to add A-overhangs. Next, the following primers were annealed to generate the splinkerette adapters: 5’-phos-CATGGTTGTTAGGACTGGAGGGGAAATCAATCCCCT-3 and 5’-CCTCCACTACGACTCACTGAAGtGCAAGCAGTCCTAACAACCATGT-3’. The adapter was ligated to the A-tailed, digested genomic DNA by O/N incubation at 16°C with T4 DNA ligase (NEB). Subsequently, ligated product was purified from the ligation reaction mixture using PCR purification, and used as template in a PCR reaction using the P-Short V2 primer (5’-CCTCCACTACGACTCACTGAAGTGC-3’) that binds the adapter and either the N-out (5’-GCGATCTAATTCTCCCCCGC-3’) or the C-out (5’-CGGGACGTCCTTCTGCTAC-3’) primers, which bind in the DSB-Spectrum cassette at the 5’-end or 3’-end, respectively. PCR amplification was done using LongAmp Taq DNA polymerase (NEB). Next, 1 μl of this first PCR reaction was used as a template for a second, nested PCR using LongAmp Taq polymerase with the primer P-nested (5’-GTGCAAGCAGTCCTAACAACCATG-3’), together with either N-in (5’-TTTGGCGTACTCACCAGTCG-3’) or C-in (5’-TCAATCCAGCGGACCTTCC-3’) primers. After amplification, the PCR reaction mixture was directly cloned into the TOPO-TA backbone (ThermoFisher Scientific) and transformed into bacteria. Plasmid DNA was extracted from bacterial colonies and sequenced. In some cases, after the second, nested PCR, the reaction mixture was analyzed by DNA gel electrophoresis, and the high abundance PCR products were gel-extracted. The extracted PCR products were subsequently amplified by repeating the second PCR, followed by TOPO-cloning and sequence analysis. All integration sites were verified by PCR.

### Statistical Analysis

All statistical analysis was performed using Graphpad Prism software. In case of direct statistical comparison between two samples a two-sided Student’s t-test was performed. In case multiple samples were tested against a control, a one-way ANOVA with post-hoc multiple comparison was performed.

## Acknowledgements

We thank the Robert A. Swanson (1969) Biotechnology Center (Koch Institute, MIT, Cambridge MA, USA) and the LUMC Flow cytometry core facility (LUMC, Leiden, the Netherlands) for assistance. This research was supported by National Institute of Health grants R01-ES015339 (M.B.Y.), R35-ES028374 (M.B.Y.), R01-CA226898 (M.B.Y.), by the Charles and Marjorie Holloway Foundation (M.B.Y.), by the MIT Center for Precision Cancer Medicine, by fellowships from the Dutch Cancer Society (BUIT 2015-7546) and the Ludwig Center at MIT’s Koch Institute (B.v.d.K.), and by ERC Consolidator (ERC-CoG-617485) and NWO-VICI grants (VI.C.182.052; H.v.A.). Support was also provided by the Cancer Center Support Grant P30-CA14051 from the National Cancer Institute and the Center for Environmental Health Sciences Support Grant P30-ES002109 from the National Institute of Environmental Health Sciences.

---

**Supplementary figure 1.**
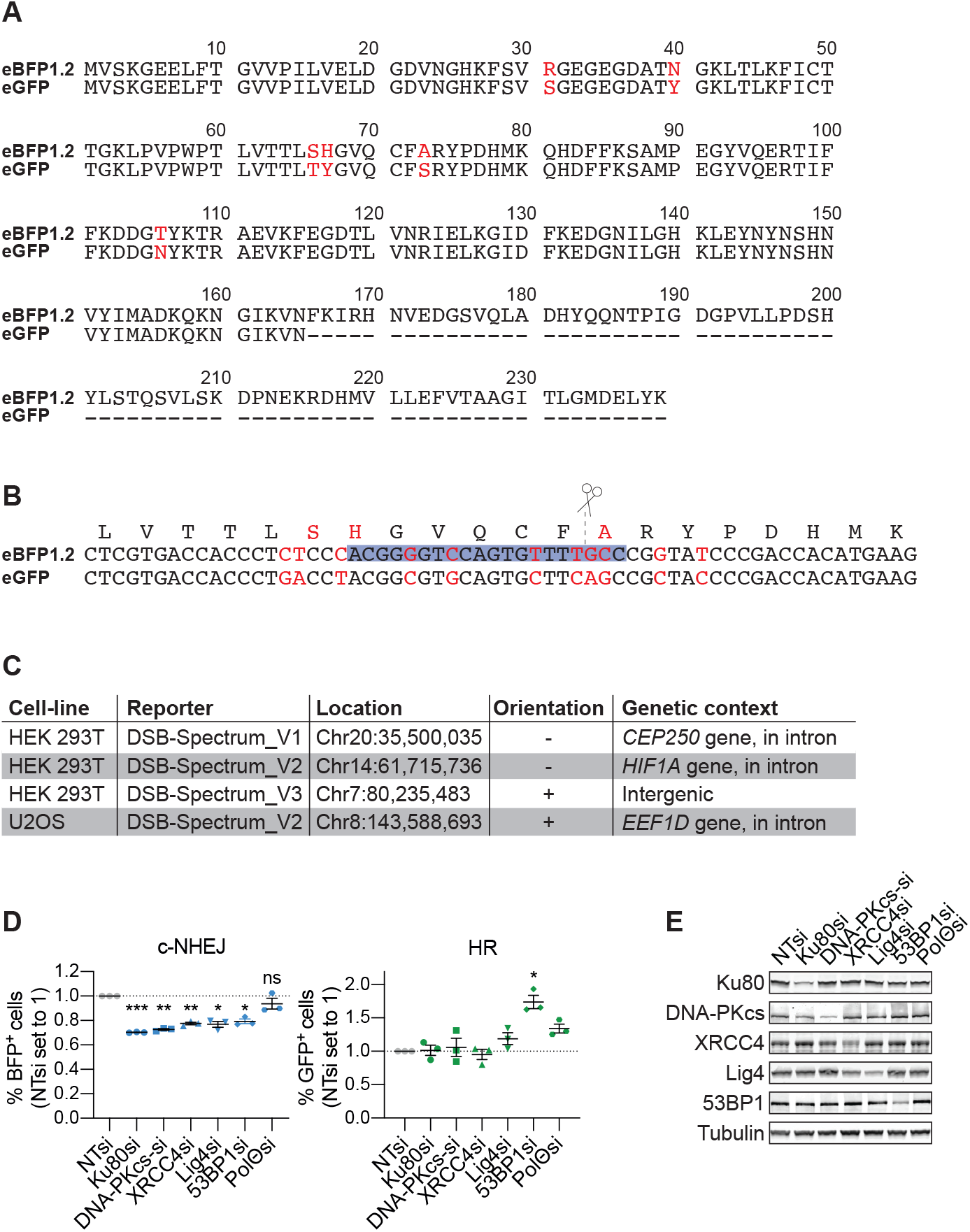
Alignment of eBFP1.2 and eGFP sequences used in DSB-Spectrum and validation of DSB-Spectrum_V1. **(A)** Alignment of the amino acid sequence translated from the eBFP1.2 and eGFP genes present in the DSB-Spectrum_V2/V3 reporter constructs. The amino acids that differ between the two sequences are indicated in red. **(B)** Alignment of the DNA sequence of the eBFP1.2 and eGFP genes in the DSB-Spectrum_V2/V3 reporter constructs. Shown is the region that contains the BFP sgRNA target site. The sgRNA sequence is indicated by the blue box. The nucleotides that differ between the two sequences are indicated in red. Note that several silent mutations were introduced in the eGFP sequence to prevent any binding of the BFP-targeting sgRNA and cutting by Cas9. **(C)** For the clonal DSB-Spectrum reporter cell-lines that were used in this manuscript, the genomic integration site of the reporter construct was determined by Splinkerette PCR as described in the materials and methods section. **(D)** DSB-Spectrum_V1 cells were transfected with indicated siRNAs, followed by transfection with Cas9 and an sgRNA targeting a control locus or BFP. At 72h after Cas9 transfection cells were analyzed by flow cytometry (n=3; mean ± SEM; ***p≤0.001; **p≤0.01; *p≤0.05; ns=non-significant). **(E)** Western blot of lysates from cells analyzed in panel D.

**Supplementary figure 2.**
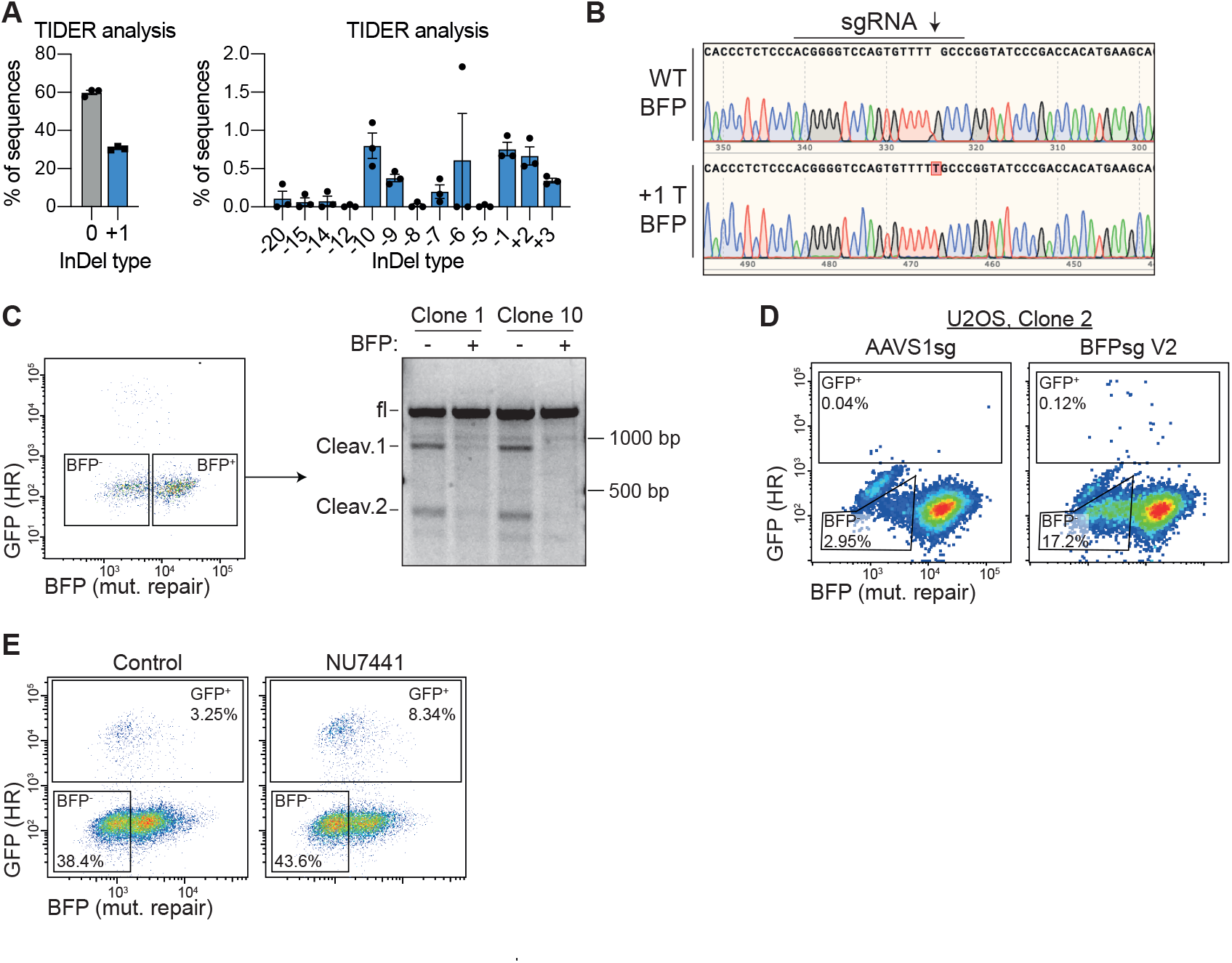
Validation of DBS-Spectrum_V2. **(A)** Replotting of the data depicted in figure 2F to show all identified InDel types. For ease of interpretation, the graph is split to show the frequency of WT and +1 sequences in the left panel, and all the other identified sequences in the right panel. **(B and C)** HEK 293T DSB-Spectrum_V2 cells were transfected with Cas9 and a BFP-targeting sgRNA, followed by FACS to collect the BFP-and BFP+ populations, and subsequent PCR amplification of the Cas9 target locus. The PCR product of the BFP-population was TOPO-cloned followed by Sanger-sequencing of individual bacterial colonies. Panel B displays a screenshot of the aligned sequencing results of a WT and +1T insertion repair product, which was the most frequent InDel identified in the BFP-population. In addition, the PCR products from both the BFP- and BFP+ populations were denatured and re-annealed, followed by Surveyor nuclease digestion and electrophoretic analysis of the digestion products, shown in panel C. Clone 1 and clone 10 refers to two different clonally derived DSB-Spectrum_V2 cell-lines. **(D)** A second U2OS DSB-Spectrum_V2 cell-line, derived from a different clone than the U2OS DSB-Spectrum_V2 cell-line shown in figure 2, was transfected with Cas9 and either a control sgRNA (AAVS1sg) or a BFP-targeting sgRNA, followed by flow cytometric analysis 72h later. Shown is a representative flow cytometry plot. **(E)** Representative flow cytometry plots of the experiments shown in figure 2I.

**Supplementary figure 3.**
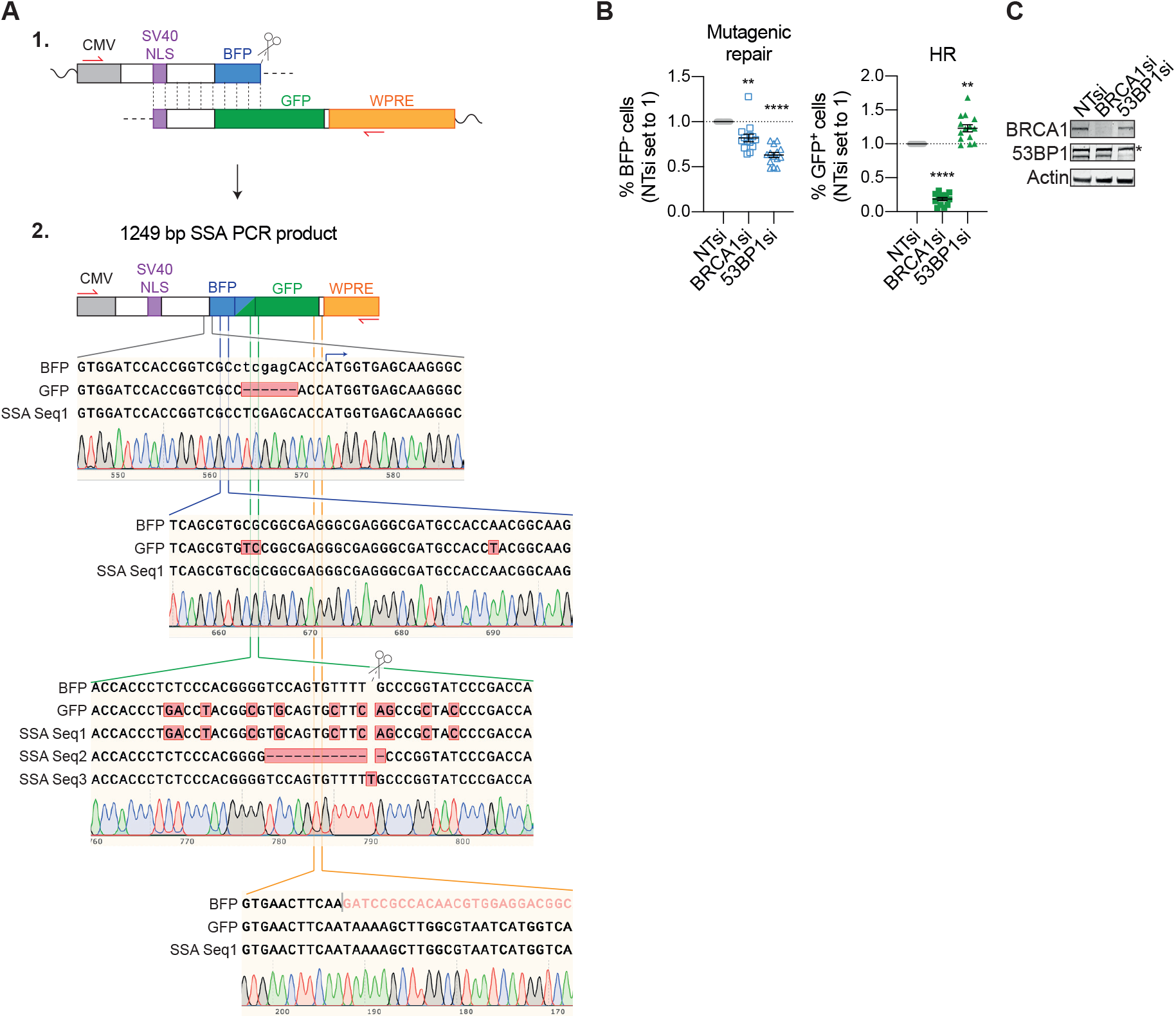
Sequence confirmation of the DSB-Spectrum_V2/V3 SSA-repair product and validation of the DBS-Spectrum_V2 reporter cells. **(A)** Diagram 1 shows a detailed schematic of the homologous regions within the DSB-spectrum reporter that are predicted to anneal during SSA-mediated repair of the Cas9-induced DSB in the BFP gene. The zigzag lines indicate omitted upstream and downstream reporter regions, the dashed lines indicate the omitted region between the homologous BFP and GFP sequences. The repair products that were amplified from the DSB-spectrum locus using the red primers (also see figure 3D) were sequence analyzed. Diagram 2 shows a schematic of the composition of the sequenced 1249 bp PCR product, validating that it is the result of DSB-repair by SSA. Shown below diagram 2 are alignments between the BFP region in DSB-Spectrum, the GFP region in DSB-Spectrum and sequences of the SSA repair product. Four alignments are shown: the top two show that the region in the SSA product located at the 5’-end of the DSB-site is a perfect match to BFP, as expected. The third panel shows an alignment of the region surrounding the Cas9-target site. As shown, in this particular region, the exact sequence of individual SSA DSB-repair products can vary. The bottom alignment shows that the region in the SSA product located at the 3’-end of the DSB-site is a perfect match to GFP, as expected. Below each alignment a representative chromatogram of the SSA repair product sequencing results is shown. **(B)** BRCA1 and 53BP1 expression was silenced by RNAi, and mutagenic repair and HR were quantified as in figure 1E but for DSB-Spectrum_V2 cells (n=14; mean ± SEM; **p≤0.01; ****p≤0.0001). **(C)** Western blot of lysates from cells analyzed in panel B. Asterisk indicates non-specific background band.

**Supplementary figure 4.**
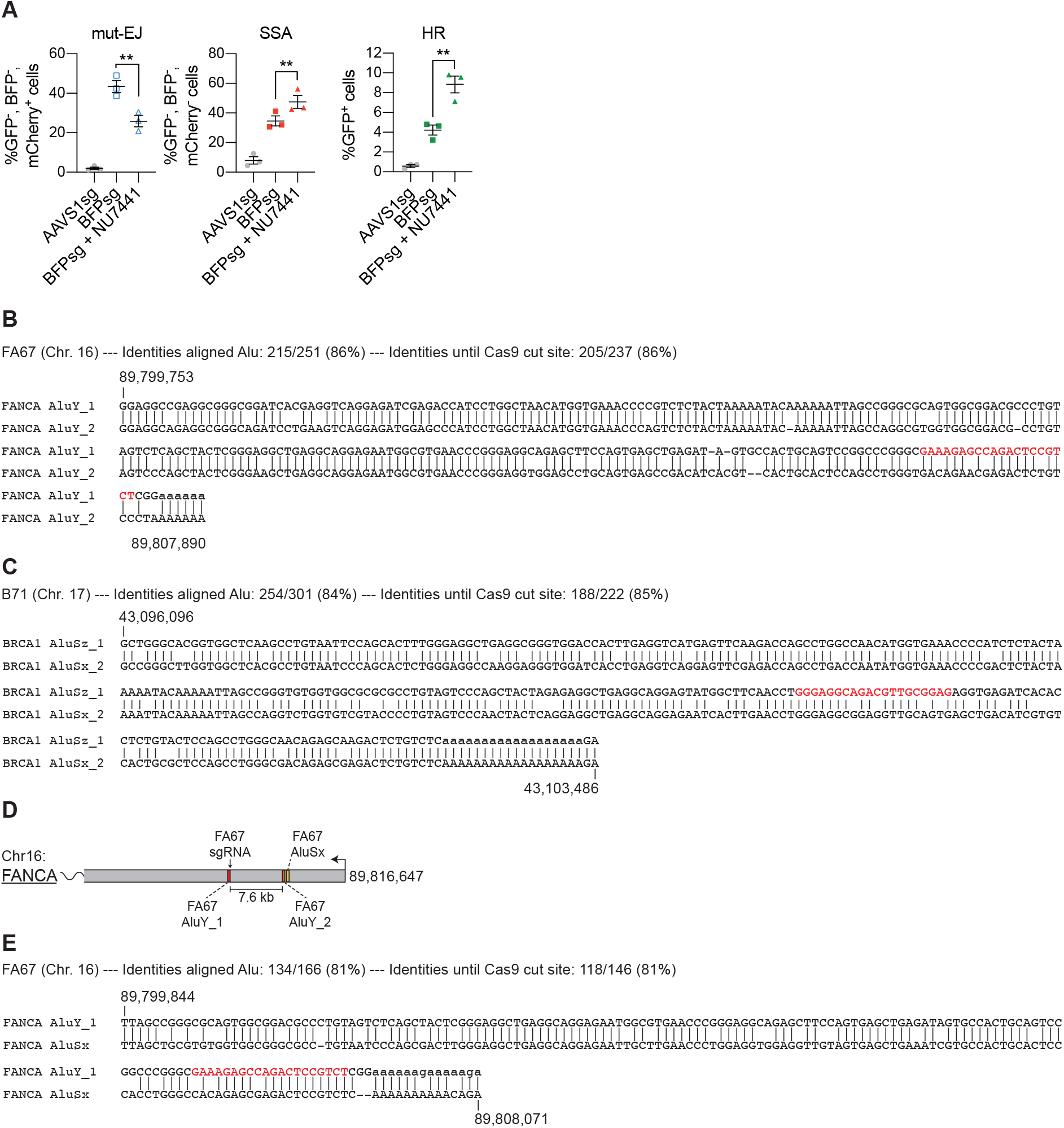
Alignment of homologous Alu elements that anneal during DSB-repair of tested Cas9 target sites. **(A)** Experiment performed as in figure 4D, in a separate HEK 293T DSB-Spectrum_V3 clone (n=3; mean ± SEM; *p≤0.05; **p≤0.01; ***p≤0.001). **(B and C)** Alignment of the homologous Alu elements that anneal during SSA-mediated DSB-repair of the FA67 (panel B) and B71 (panel C) Cas9 target sites in the *FANCA* and *BRCA1* genes respectively (also see figure 5). The chromosomal location (according to human genome assembly hg38) of the first aligned nucleotide of the upstream and the last aligned nucleotide of the downstream Alu-element are indicated on the top left and bottom right of the alignment, respectively. The sgRNA target sites are indicated in red. **(D)** As in figure 5E, but including the AluSx element (yellow box) that is located directly downstream of the AluY_2 element. **(E)** As in panel B, but now aligning the homologous AluY_1 and AluSx elements.

**Supplementary figure 5.**
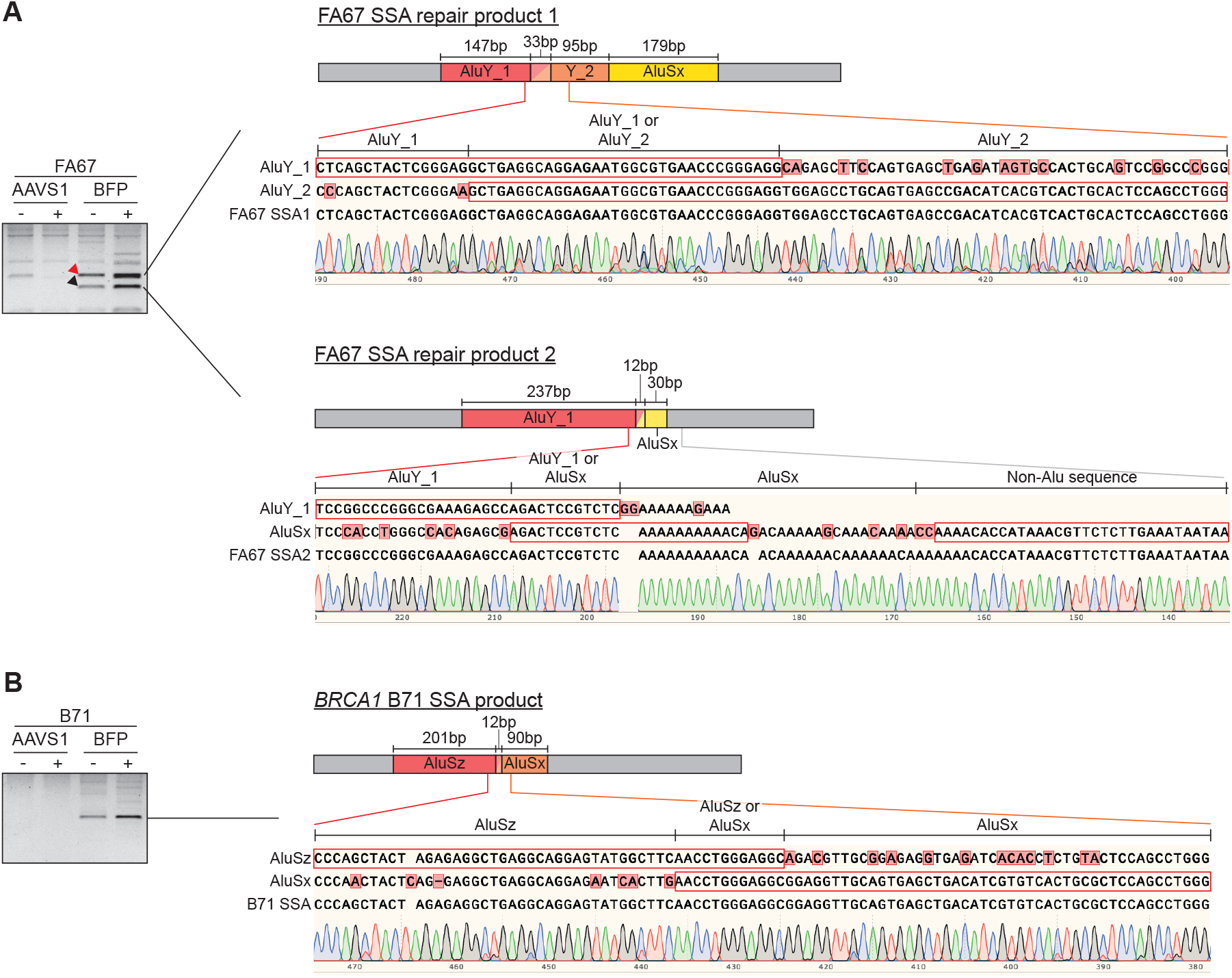
Sequence validation of the FA67 and B71 SSA DSB-repair products. **(A and B)** The FA67 and B71 SSA repair products that were PCR amplified as described in figure 5 were TOPO-cloned and analyzed by Sanger sequencing. DNA gel electrophoresis results shown in figure 5I are depicted again here to indicate which PCR products were sequence-analyzed. The composition of the SSA repair products were determined based on this sequence analysis and displayed as cartoons indicating the length of each of the remaining Alu elements. Shown below these cartoons is an alignment of the sequence of the individual Alu elements to the sequence of the SSA repair product, focussing on the junction region between the two recombined Alu elements. Regions in each Alu element that align perfectly to the SSA repair product are indicated by the box. Positions in the Alu element sequence that are mismatches to the SSA repair product sequence are indicated by red shading. Below the alignment a representative chromatogram of the SSA repair product sequencing results is shown. Note that regions of the Alu elements can share 100% homology, and it is therefore not possible to assign in the SSA repair product whether these regions are derived from the upstream or downstream Alu element. For example, in the FA67 SSA repair product 1, there is a 33 bp region at the junction between the AluY_1 and AluY_2 element that could be derived from either element.

**Supplementary figure 6.**
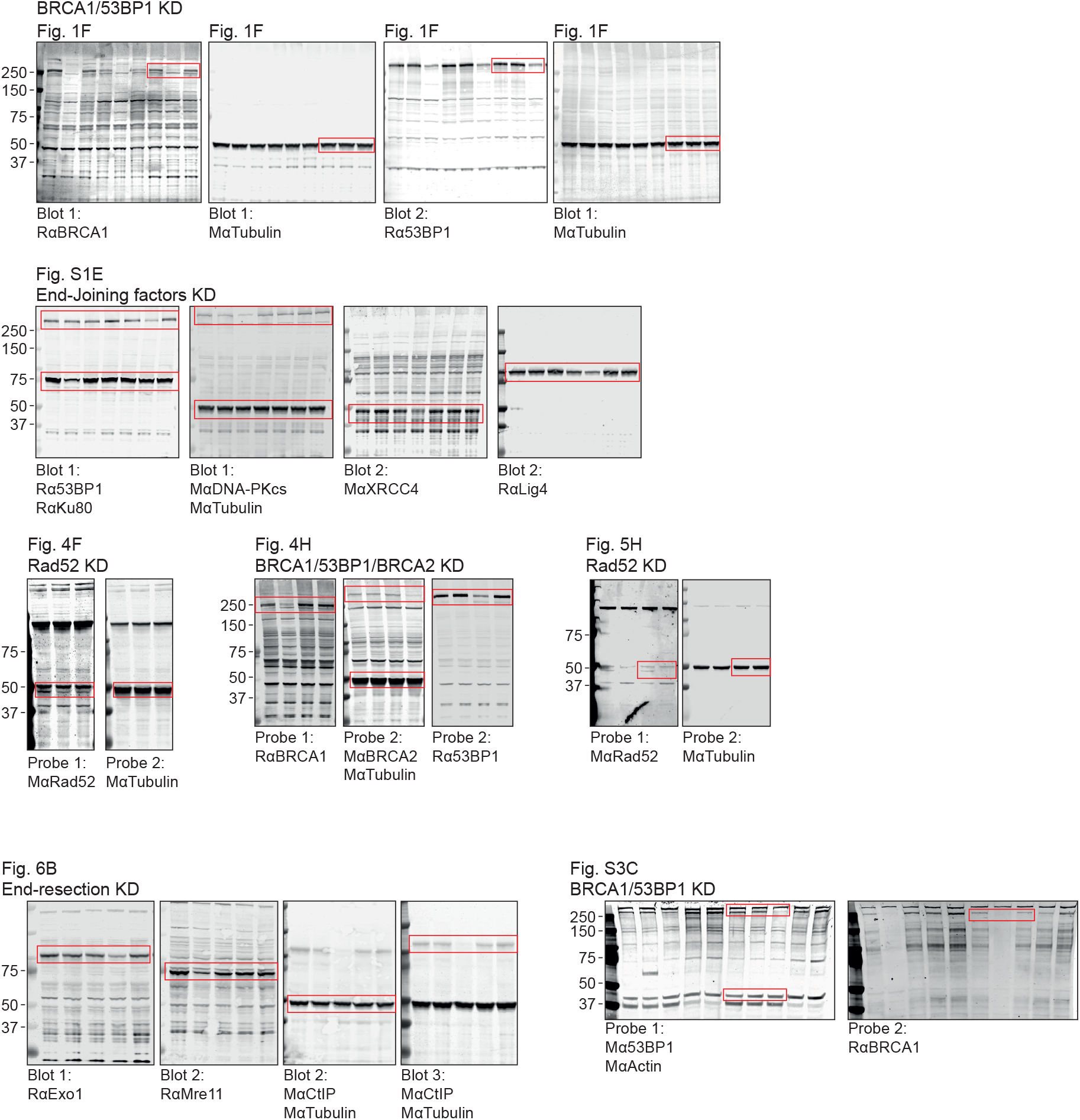
Uncropped western blot images for all indicated figure panels. Red boxes indicate cropped areas displayed in annotated figure panels.

## References

1. Cannan, W. J. & Pederson, D. S. Mechanisms and Consequences of Double-Strand DNA Break Formation in Chromatin. J. Cell. Physiol. 231, 3–14 (2016).

2. Komor, A. C., Badran, A. H. & Liu, D. R. CRISPR-Based Technologies for the Manipulation of Eukaryotic Genomes. Cell 168, 20–36 (2017).

3. Doudna, J. A. The promise and challenge of therapeutic genome editing. Nature 578, 229–236 (2020).

4. Knott, G. J. & Doudna, J. A. CRISPR-Cas guides the future of genetic engineering. Science. 361, 866–869 (2018).

5. Scully, R., Panday, A., Elango, R. & Willis, N. A. DNA double-strand break repair-pathway choice in somatic mammalian cells. Nat. Rev. Mol. Cell Biol. 20, 698–714 (2019).

6. Pannunzio, N. R., Watanabe, G. & Lieber, M. R. Nonhomologous DNA end-joining for repair of DNA double-strand breaks. J. Biol. Chem. 293, 10512–10523 (2018).

7. Jasin, M. & Rothstein, R. Repair of Strand Breaks by Homologous Recombination. Cold Spring Harb. Perspect. Biol. 5, (2013).

8. Yeh, C. D., Richardson, C. D. & Corn, J. E. Advances in genome editing through control of DNA repair pathways. Nat. Cell Biol. 21, 1468–1478 (2019).

9. Bhargava, R., Onyango, D. O. & Stark, J. M. Regulation of Single-Strand Annealing and its Role in Genome Maintenance. Trends Genet. 32, 566–575 (2016).

10. Sallmyr, A. & Tomkinson, A. E. Repair of DNA double-strand breaks by mammalian alternative end-joining pathways. J. Biol. Chem. 293, 10536–10546 (2018).

11. Kelso, A. A., Lopezcolorado, F. W., Bhargava, R. & Stark, J. M. Distinct roles of RAD52 and POLQ in chromosomal break repair and replication stress response. PLOS Genet. 15, e1008319 (2019).

12. Mendez-Dorantes, C., Bhargava, R. & Stark, J. M. Repeat-mediated deletions can be induced by a chromosomal break far from a repeat, but multiple pathways suppress such rearrangements. Genes Dev. 32, 524–536 (2018).

13. Shen, M. W. et al. Predictable and precise template-free CRISPR editing of pathogenic variants. Nature 563, 646–651 (2018).

14. Allen, F. et al. Predicting the mutations generated by repair of Cas9-induced double-strand breaks. Nat. Biotechnol. 37, 64–72 (2019).

15. Knijnenburg, T. A. et al. Genomic and Molecular Landscape of DNA Damage Repair Deficiency across The Cancer Genome Atlas. Cell Rep. (2018).

16. Zimmermann, M., Lottersberger, F., Buonomo, S. B., Sfeir, A. & De Lange, T. 53BP1 regulates DSB repair using Rif1 to control 5′ end resection. Science. 339, 700–704 (2013).

17. Chapman, J. R. et al. RIF1 Is Essential for 53BP1-Dependent Nonhomologous End Joining and Suppression of DNA Double-Strand Break Resection. Mol. Cell 49, 858–871 (2013).

18. Escribano-Díaz, C. et al. A Cell Cycle-Dependent Regulatory Circuit Composed of 53BP1-RIF1 and BRCA1-CtIP Controls DNA Repair Pathway Choice. Mol. Cell 49, 872–883 (2013).

19. Noordermeer, S. M. et al. The shieldin complex mediates 53BP1-dependent DNA repair. Nature 560, 117–121 (2018).

20. Gupta, R. et al. DNA Repair Network Analysis Reveals Shieldin as a Key Regulator of NHEJ and PARP Inhibitor Sensitivity. Cell 173, 972–988 (2018).

21. Dev, H. et al. Shieldin complex promotes DNA end-joining and counters homologous recombination in BRCA1-null cells. Nat. Cell Biol. 20, 954–965 (2018).

22. Trenner, A. & Sartori, A. A. Harnessing DNA Double-Strand Break Repair for Cancer Treatment. Frontiers in Oncology 9, 1388 (2019).

23. Pierce, A. J., Johnson, R. D., Thompson, L. H. & Jasin, M. XRCC3 promotes homology-directed repair of DNA damage in mammalian cells. Genes Dev. 13, 2633–2638 (1999).

24. Stark, J. M., Pierce, A. J., Oh, J., Pastink, A. & Jasin, M. Genetic Steps of Mammalian Homologous Repair with Distinct Mutagenic Consequences. Mol. Cell. Biol. 24, 9305–9316 (2004).

25. Bhargava, R. et al. C-NHEJ without indels is robust and requires synergistic function of distinct XLF domains. Nat. Commun. 9, 2484 (2018).

26. Bennardo, N., Cheng, A., Huang, N. & Stark, J. M. Alternative-NHEJ is a mechanistically distinct pathway of mammalian chromosome break repair. PLoS Genet. 4, e1000110 (2008).

27. Kuhar, R. et al. Novel fluorescent genome editing reporters for monitoring DNA repair pathway utilization at endonuclease-induced breaks. Nucleic Acids Res. 42, e4–e4 (2014).

28. Certo, M. T. et al. Tracking genome engineering outcome at individual DNA breakpoints. Nat. Methods 8, 671–676 (2011).

29. Yin, B. & Largaespada, D. A. PCR-based procedures to isolate insertion sites of DNA elements. Biotechniques 43, 79–84 (2007).

30. Leahy, J. J. J. et al. Identification of a highly potent and selective DNA-dependent protein kinase (DNA-PK) inhibitor (NU7441) by screening of chromenone libraries. Bioorg. Med. Chem. Lett. 14, 6083–6087 (2004).

31. Brinkman, E. K., Chen, T., Amendola, M. & van Steensel, B. Easy quantitative assessment of genome editing by sequence trace decomposition. Nucleic Acids Res. 42, e168–e168 (2014).

32. Brinkman, E. K. et al. Easy quantification of template-directed CRISPR/Cas9 editing. Nucleic Acids Res. 46, e58–e58 (2018).

33. van Overbeek, M. et al. DNA Repair Profiling Reveals Nonrandom Outcomes at Cas9-Mediated Breaks. Mol. Cell 63, 633–646 (2016).

34. Qiu, P. et al. Mutation detection using Surveyor™nuclease. Biotechniques 36, 702–707 (2004).

35. Chen, C.-C., Feng, W., Lim, P. X., Kass, E. M. & Jasin, M. Homology-Directed Repair and the Role of BRCA1, BRCA2, and Related Proteins in Genome Integrity and Cancer. Annu. Rev. Cancer Biol. 2, 313–336 (2018).

36. Batzer, M. A. & Deininger, P. L. Alu repeats and human genomic diversity. Nat. Rev. Genet. 3, 370–379 (2002).

37. Riesenberg, S. et al. Simultaneous precise editing of multiple genes in human cells. Nucleic Acids Res. 47, e116–e116 (2019).

38. Robert, F., Barbeau, M., Éthier, S., Dostie, J. & Pelletier, J. Pharmacological inhibition of DNA-PK stimulates Cas9-mediated genome editing. Genome Med. 7, 93 (2015).

39. Ochs, F. et al. 53BP1 fosters fidelity of homology-directed DNA repair. Nat. Struct. Mol. Biol. 23, 714–721 (2016).

40. Gunn, A. & Stark, J. M. I-SceI-based assays to examine distinct repair outcomes of mammalian chromosomal double strand breaks. Methods Mol. Biol. 920, 379–391 (2012).

41. Gomez-Cabello, D., Jimeno, S., Fernández-Ávila, M. J. & Huertas, P. New Tools to Study DNA Double-Strand Break Repair Pathway Choice. PLoS One 8, e77206 (2013).

42. Glaser, A., McColl, B. & Vadolas, J. GFP to BFP Conversion: A Versatile Assay for the Quantification of CRISPR/Cas9-mediated Genome Editing. Mol. Ther. - Nucleic Acids 5, e334 (2016).

43. Roidos, P. et al. A scalable CRISPR/Cas9-based fluorescent reporter assay to study DNA double-strand break repair choice. Nat. Commun. 11, 4077 (2020).

44. Tennant, P. A. et al. Fluorescent in vivo editing reporter (FIVER): A novel multispectral reporter of in vivo genome editing. bioRxiv 2020.07.14.200170 (2020).

45. Eki, R. et al. A robust CRISPR–Cas9-based fluorescent reporter assay for the detection and quantification of DNA double-strand break repair. Nucleic Acids Res. (2020).

46. Liang, F., Han, M., Romanienko, P. J. & Jasin, M. Homology-directed repair is a major double-strand break repair pathway in mammalian cells. Proc. Natl. Acad. Sci. 95, 5172 LP – 5177 (1998).

47. Lander, E. S. et al. Initial sequencing and analysis of the human genome. Nature 409, 860–921 (2001).

48. Flynn, E. K. et al. Comprehensive Analysis of Pathogenic Deletion Variants in Fanconi Anemia Genes. Hum. Mutat. 35, 1342–1353 (2014).

49. Petrij-Bosch, A. et al. BRCA1 genomic deletions are major founder mutations in Dutch breast cancer patients. Nat. Genet. 17, 341–345 (1997).

50. Rohlfs, E. M. et al. An Alu-mediated 7.1 kb deletion of BRCA1 exons 8 and 9 in breast and ovarian cancer families that results in alternative splicing of exon 10. Genes, Chromosom. Cancer 28, 300–307 (2000).

51. Lord, C. J. & Ashworth, A. BRCAness revisited. Nat. Rev. Cancer (2016). doi:10.1038/nrc.2015.21

52. Gunn, A., Bennardo, N., Cheng, A. & Stark, J. M. Correct end use during end joining of multiple chromosomal double strand breaks is influenced by repair protein RAD50, DNA-dependent protein kinase DNA-PKcs, and transcription context. J. Biol. Chem. 286, 42470–42482 (2011).

53. Adikusuma, F. et al. Large deletions induced by Cas9 cleavage. Nature 560, E8–E9 (2018).

54. Kosicki, M., Tomberg, K. & Bradley, A. Repair of double-strand breaks induced by CRISPR–Cas9 leads to large deletions and complex rearrangements. Nat. Biotechnol. 36, 765–771 (2018).

55. Alanis-Lobato, G. et al. Frequent loss-of-heterozygosity in CRISPR-Cas9-edited early human embryos. bioRxiv 2020.06.05.135913 (2020).

56. Garcia, V., Phelps, S. E. L., Gray, S. & Neale, M. J. Bidirectional resection of DNA double-strand breaks by Mre11 and Exo1. Nature 479, 241–244 (2011).

57. Ronato, D. A. et al. Limiting the DNA Double-Strand Break Resectosome for Genome Protection. Trends Biochem. Sci. 45, 779–793 (2020).

58. Karanja, K. K., Cox, S. W., Duxin, J. P., Stewart, S. A. & Campbell, J. L. DNA2 and EXO1 in replication-coupled, homology-directed repair and in the interplay between HDR and the FA/BRCA network. Cell Cycle 11, 3983–3996 (2012).

59. Tomimatsu, N. et al. DNA-damage-induced degradation of EXO1 exonuclease limits DNA end resection to ensure accurate DNA repair. J. Biol. Chem. 292, 10779–10790 (2017).

60. Chen, C.-C. et al. EXO1 suppresses double-strand break induced homologous recombination between diverged sequences in mammalian cells. DNA Repair (Amst). 57, 98–106 (2017).

61. Ran, F. A. et al. Genome engineering using the CRISPR-Cas9 system. Nat. Protoc. 8, 2281–2308 (2013).

62. de Jager, M. et al. DNA-binding and strand-annealing activities of human Mre11: implications for its roles in DNA double-strand break repair pathways. Nucleic Acids Res. 29, 1317–1325 (2001).

